# Trait evolution linked to climatic shifts contributes to adaptive divergence in an alpine carnation (*Dianthus sylvestris*)

**DOI:** 10.1101/2024.03.22.586106

**Authors:** Aksel Pålsson, Simone Fior, Alex Widmer

## Abstract

Populations expanding to new habitats may encounter novel selection regimes which can lead to ecotype formation. In the Alps, elevation corresponds to steep ecological gradients, along which ecotype formation has occurred in many species. The majority of alpine plant species are perennial and little is known about how selection acts across different stages of their life-cycles and how fitness trade-offs shape adaptive processes in perennials. We investigated how selection at opposite ends of elevational gradients has driven ecotype formation in *Dianthus sylvestris*, a perennial herb that expanded its ecological niche to low elevation habitats after the Last Glacial Maximum. Through a multi-year reciprocal transplant experiment including parental populations and recombinant crosses we assessed fitness under natural conditions and dissect how adaptation is mediated by different fitness components with inherent trade-offs, and pinpoint the contribution of growth and reproductive traits to this process. We show that the evolution of local adaptation proceeded by selection acting primarily through reproduction and survival at low and high elevation, respectively. At low elevation the primary contribution to adaptation was third year reproduction, concomitant with a left skewed age distribution. At high elevation the contribution to adaptation and the age distribution were more dispersed across the life cycle. We found that large, early flowering plants have a consistent fitness advantage. This was mediated by direct selection favoring large size through reproductive output at low elevation, and early flowering through the probability to produce seeds at high elevation. Our results indicate that the selection regime imposed by the warm low elevation habitat led to the evolution of an ecotype exhibiting a life-history strategy characterized by high investment in rapid growth and early reproduction. In contrast, the high elevation strategy favors high investment in self-maintenance. Our results suggest that weakening of a key fitness trade-off associated with resource allocation contributed to the evolution of distinct ecotypes in this perennial plant species.

## Introduction

Populations facing novel environments may experience selection regimes that favor traits and trait combinations that differ from those favored in the original habitats. For a large number of species in the northern hemisphere, current distributions result from demographic responses to climatic changes following the Last Glacial Maximum (Davis & Shaw, 2001; Petit *et al.,* 2003). As environments suitable to species persistence opened up beyond glacial refugial ranges upon the retreat of the glaciers, populations expanding into deglaciated areas either tracked their ecological niche or expanded their niche into warmer habitats. The response to novel selection regimes has likely driven the formation of warm adapted ecotypes (Hargreaves *et al.,* 2014). In temperate mountain ranges, recolonization of elevational gradients exposed populations to strong clinal variation in key abiotic factors that frequently assert strong selection pressures (Körner, 2003; Keller *et al.,* 2013; Halbritter *et al.,* 2018). This has resulted in local adaptation in a large number of species, accompanied by changes in both phenotypic and life history traits (Halbritter *et al.,* 2018). Dissecting the underlying processes offers an excellent opportunity to gain insights into the evolution of contemporary ecotypes in response to climate-driven selection.

Elevational gradients form steep ecological gradients that are primarily shaped by changes in physical parameters. The shift to warmer temperatures at the onset of the present interglacial period and the retreat of the glaciers have facilitated the formation and colonization of novel, warmer habitats. Today, rapid anthropogenic climate change is predicted to bear major impacts on species’ ecological niches and distribution ranges (Tito *et al.,* 2020; Pörtner *et al.,* 2022). In most mountain systems, temperature constitutes a primary determinant of the abiotic environment, with cascading effects on multiple selective agents (Körner, 2003). Typically, increased elevation corresponds to a decrease in temperature, and lower temperatures at high elevation imply a longer period of snow cover, with consequently shorter summer seasons (Körner, 2003). This can directly slow down organisms’ metabolic processes and affect their physiology (Körner, 2006; Poorter *et al.,* 2011). Temperature further impacts biotic interactions, such as e.g., between plants and pollinators or hosts and their parasites, which are fundamental for species persistence (López-Goldar & Agrawal, 2021).

In plants, elevational ecotypes commonly exhibit phenotypic divergence with a strong genetic basis, and shared trends in fitness-related traits are observed across a diversity of taxa. In a meta-analysis on ecological evidence of elevation adaptation, Halbritter *et al*., (2018) found that elevational ecotypes display pronounced divergence in flowering time and size, expressed both as plant height and biomass. Both plant height and flowering time are essential for successful reproduction in many plant species, thus bearing direct effects on individual fitness (Gervasi & Schiestl, 2017; Gaudinier & Blackman, 2019). While selection on these traits is typically mediated by interactions with other biotic agents such as pollinators (Zu & Schiestl, 2017), climate directly impacts the fine-tuning of this crucial life stage by governing the plants’ ability to achieve physiological thresholds (Amasino, 2010; Cho *et al.,* 2017; Ehrlén *et al*., 2020). Most plants must reach a minimum size before they can reproduce (Weiner *et al.,* 2009; Younginger *et al*., 2017), and individuals with a higher ability to accumulate biomass have more resources to invest in reproductive structures (Bonser & Aarssen, 2009; Cheplick, 2020; Proulx, 2021). Because size shares a positive allometric relationship with temperature, this physical parameter affects the evolution of reproductive traits and of life-history strategies that optimize the antagonistic resource requirements for self-maintenance and reproduction (Stearns, 1992). This is reflected in vegetation at the warmer end of elevational gradients that often consists of plants exhibiting a life-history strategy characterized by large, fast-growing plants with high investment in reproduction, whereas at the opposite end, populations typically display a more conservative strategy, characterized by slower growth and a longer lifespan (von Arx *et al.,* 2006; Hautier *et al.,* 2009; Kim & Donohue, 2011; Kim & Donohue, 2013; Laiolo & Obeso, 2017; Rosbakh & Poschlod, 2018). While the phenotypic variation in plant size, height and phenology and the consequential divergence in life-history traits along elevational gradients is well documented, the mechanisms by which these traits interact in their contribution to adaptation remain largely unknown.

Unravelling the evolution of phenotypic traits in different environments requires to assess how variation in trait values expressed by different individuals correlates with relative fitness. Heritable trait variation that is consistently associated with fitness may drive trait divergence between environments, thus resulting in phenotypic divergence between locally adapted ecotypes. Fitness itself results from the combined effects of separate components, such as survival and reproduction (Orr, 2009; Acerenza, 2016), which may act coherently in favoring optimal trait values, but also drive contrasting trajectories (Wadgymar *et al*., 2017). In *Mimulus guttatus*, for example, selection favors larger flowers through reproduction while simultaneously favoring smaller flowers through survivorship, whose stronger effect during the entire life cycle eventually results in smaller flowers being advantageous (Mojica & Kelly, 2010). These complex trait-fitness interactions complicate the study of adaptation, and call for the identification of fitness components with a major role in ensuring population persistence. Moreover, as the relative contribution of separate fitness components may vary depending on the environment, selection may act through alternative fitness components in different ecotypes (Goebl *et al*., 2022). Hence, understanding the role of phenotypic traits in adaptation requires experiments to dissect the process of phenotypic selection though both individual and combined analyses of multiple fitness components.

Studies of local adaptation are ideally performed using reciprocal transplant experiments where genotype by environment interactions can be tested (Johnson *et al*., 2021). In such experiments, populations are reciprocally transplanted between habitats and local adaptation is inferred through two criteria (Kawecki & Ebert, 2004). The native populations display higher fitness in their native habitat relative to the foreign population (local vs. foreign criterion) and they have higher fitness in their native habitat than when growing in the foreign habitat (home vs. away criterion). The first of these criteria is generally regarded as a stronger indicator of local adaptation. While ecotypes constitute natural units to test for local adaptation, they are of limited utility to dissect the role of individual traits in this process. In locally adapted populations, selection has driven the evolution of different phenotypes that maximize fitness, so that individuals express correlated trait values, with limited variation centered around optimum values (Lexer *et al*., 2003; Ferris & Willis, 2018). Phenotypic selection analyses, however, require substantial trait variation to assess its effects on fitness. This can be achieved by exposing genotypes from experimental crosses to natural selection, where reshuffling of the genomes through recombination can generate a wider distribution of phenotypic values than those expressed in local ecotypes.

*Dianthus sylvestris* (Caryophyllaceae) is a widespread perennial herb that occupies elevational gradients from the colline to the alpine belt of the European Alps (Collin & Shykoff, 2003; Info Flora 2016). Previous work has shown that during the Last Glacial Maximum, D. sylvestris survived in south-eastern refugia characterized by narrow climatic conditions similar to those of present day high elevation habitats (Luqman *et al*., 2023). Post-glacial warming has then facilitated recolonization of the Alpine Arch concomitant with the expansion of the species’ climatic niche through adaptive evolution in the warm habitats at low elevation. Consequently, contemporary elevational ecotypes show substantial divergence in fitness-related traits consistent with the physiological response to climate driven selection of many alpine species, in particular in plant size, plant height and flowering time. The derived condition of these traits in *D. sylvestris* offers an excellent opportunity to study the evolutionary trajectories underlying the species’ response to recent climate driven selection.

In this study, we use ecologically diverged elevational populations of *D. sylvestris* from the central Alps to dissect how selection acting through alternate fitness components has driven ecotype formation and assess the contribution of phenotypic divergence in plant size, plant height and flowering time to this process. We use a transplant experiment of wild populations from the extremes of the elevational gradient occupied by the species to find evidence of adaptation and complement our field trials with recombinant populations to perform phenotypic selection analyses and dissect the adaptive role of fitness-related traits. Specifically, we ask; 1) Does the phenotypic divergence of plant size, plant height and flowering time between populations inhabiting the high elevation and low elevation habitats have a genetic basis? 2) Has the colonization of the low elevation habitat driven the evolution of elevational adaptation? 3) If so, is this concomitant with variation in life history traits favoring alternative strategies of resource allocation to self-maintenance vs. reproduction? 4) How does the phenotypic divergence in plant size, plant height and flowering time between the elevational ecotypes contribute to the recent adaptation to a warmer environment?

## Methods

### Experimental set up

#### Reciprocal transplantation of wild populations

We sampled seeds from 13 to 41 individuals in wild populations of *D. sylvestris* at three low– (i.e., <1000 m.a.s.l.) and three high– (i.e., > 2000 m.a.s.l.) elevation sites in the central Alps (Valais, Switzerland; Figure S1, Table S1). During summer 2015, seeds were germinated in a greenhouse (Lindau-Eschikon, Switzerland) in peat moss based soil (Klasmann Deilmann Gmbh) under a 12-hour day/night cycle at 20/18°C and relative humidity of 50-60%. In fall 2015, seedlings were transferred to our transplant sites in geographic proximity to the six source populations (Pålsson *et al.,* 2023b). Climatic conditions at our two high and two low elevation sites resemble those of our six natural populations, in particular in respect to temperature and precipitation (Pålsson *et al.,* 2023b). We transplanted ∼300 seedlings into each of four transplant sites, providing even representation of populations and seed families across the four sites. Seeds were arranged randomly in 25 cm grids in 6 blocks of ∼72 individuals each. Blocks further included 3-22 F1 individuals that were not used in this study and were randomly placed within each site, along with 8 blocks of *Dianthus carthusianorum* that were also not used here. We fenced transplant sites to exclude mammalian herbivores and regularly trimmed the vegetation surrounding our plants to prevent our plants from being overgrown. Hence, our experiments primarily test responses to abiotic factors, although biotic components such as below-ground competition or plant-pollinator interactions are integral part of the observed natural processes. To ensure successful establishment, we watered the plants twice a week during the first winter of 2015/2016. Plants that died during this period were attributed to transplant shock and excluded from further analyses. At the start of 2016, 690 and 645 plants were alive in the low and high elevation sites, respectively, and form the basis of our analyses (Table S2).

#### Seedling recruitment experiment

In fall 2020, we sowed seeds produced in 2018 in each of our transplant sites to obtain an in-situ estimate of seedling recruitment. Whenever possible, we pooled 100 seeds per population equally representing five maternal families and sowed five seeds per family in each of 20 biodegradable pots (approximately: 10×8×6 cm, 0.4 litres) containing peat moss soil (Klasmann Deilmann Gmbh) (Table S3). Due to variation in seed production among populations and sites, in some cases the number of seeds sown per population and seed family varied (Table S3). We embedded the pots in positions within the blocks of the original experiment. Seedlings alive at the end of 2021 were considered successful recruitments.

#### Transplantation of recombinant populations

We generated recombinant (F2) plants derived from two F1 families that were derived from controlled crosses between one low– and one high-elevation individual (Table S4). During the flowering season 2017, we placed 15-28 F1 individuals in each of five cages at the research station Lindau-Eschikon and weekly supplemented butterflies (*Pieris brassicae*) to act as pollinators. With this approach, three types of recombinant groups are potentially produced in each cage: descendants of each of the two F1 lines, and hybrids between these. Seeds were then germinated as reported above and ∼1000 seedlings representative of the three types of crosses were transplanted into 14 new blocks each added to one low– and one high-elevation site with the same grid design as described above. In spring 2018, 621 and 555 plants survived in the low– and high-elevation site, respectively (Table S5).

#### Data collection

We collected data on flowering time (i.e., date of anthesis of the first flower), plant height, plant size, and fitness over five (2016 – 2020) and three (2018 – 2020) growing seasons for the wild and the F2 populations, respectively. The growing season typically spans March to November in the low sites and May to October in the high sites (for details see, Pålsson *et al.,* 2023b). At the start and end of each growing season, we took high resolution images (Nikon D810; 7360×4912 pixels) of all individual plants, scored survival and estimated plant size as the mean of two orthogonal diameters of the rosettes measured using image J v.2.0 (Schindelin *et al.,* 2012). During the growing seasons, we visited each site twice a week in 2016 and once a week in the following years. For the recombinant populations, no flowering occurred in 2018, and we visited sites twice a week during the 2019 and 2020 growing seasons. After wilting of the flowers, we bagged inflorescences in organza bags and harvested all stalks when the seeds had ripened. Plants typically produce stalks of similar length, and we estimated plant height as the mean length of three representative stalks. In the laboratory we extracted seeds from capsules and estimated individual seed output for the wild populations as the average number of seeds from two independent runs of an elmor C3 High Sensitive Seed Counter (elmor Ltd, Schwyz, Switzerland). For F2 individuals, we weighed all seeds produced per plant using a Mettler Toledo Ae240 Balance (precise to the nearest 0.0001 gram). For analyses requiring count data, cumulative seed weight was transformed to seed number using correlations inferred from a subset of ∼80 individuals from the low– and high-elevation site (Figure S2).

## Statistical analyses

In all analyses of the wild populations, we considered the low and high elevation populations as representative of the low and high elevation ecotypes, and transplant sites at the same elevation as replicates of the low and high elevation environments. All analyses were performed in R v. 3.3.2 (R Core Development Team 2016).

### Phenotypic divergence between elevational ecotypes

We tested for phenotypic divergence between elevational ecotypes in flowering time, plant height and plant size, both in mean treat expression and at subsequent time points across the life cycle. We standardized the traits within each year and elevational environment to a mean of 0 and standard deviation of 1. Traits were set as response variable in linear mixed effect models (LMMs) with Gaussian error distributions and ecotype, transplant environment and their ecotype by environment interaction set as predictor variables. Trait values were log transformed when this improved distribution of the residuals (see results). To uncover genetic correlations between plant size, plant height and flowering time, we fitted the latter two as response variables and plant size, ecotype, and transplant environment and their interaction as predictor variables in LMMs. In all analyses, we nested seed families within population and block nested within site as random effects. These were excluded in a few cases when low variation caused singular fits of the models (see results).

### Elevational adaptation

To test for elevational adaption of the two ecotypes, we formulated age-structured matrix population models (MPM) yielding an integrated estimate of fitness expressed as population growth rate (λ), as described in Pålsson *et al.,* (2023b). Briefly, we divided the life cycle into winter (W_i_) and summer (S_i_) stages and calculated the survival vital rates (T_i_) as the proportion of individuals transitioning across stages (Figure S3) (Caswell, 2001). Reproductive vital rates (R_i_) were estimated as the product of flowering probability, seed count and recruitment. Our empirical recruitment estimates recovered overall higher estimates for both ecotypes in the low (i.e., low ecotype: 0.19 ± 0.19; high ecotype 0.16 ± 0.20) compared to high environment (i.e., low ecotype: 0.06 ± 0.06; high ecotype: 0.13 ± 0.16). To assess whether differences in population growth rates of ecotypes growing in each environment are statistically significant, we performed 20 000 bootstrap replicates of each matrix stratified by population, and estimated bias corrected 95% confidence intervals around all means. These analyses were performed in the packages popbio v. 2.2.4 (Stubben & Milligan, 2007) and boot v. 1.3 (Canty & Ripley, 2021). To further investigate the contribution of individual fitness components to adaptation in the transplant environments, we decomposed the contributions of specific vital rates to variation in λ by using life-table response experiments (LTRE) (Caswell, 1989). These tested the matrix of the foreign ecotype against the matrix of the native ecotype in each environment. LTRE contributions were inferred from 20 000 bootstrap replicates of each matrix stratified by population.

### Contributions of single fitness components to adaptation

We dissected the contributions of three separate fitness components to adaptation: flowering probability, survival probability and seed count. For each fitness component, we tested ecotype x environment interactions, and for differential performance of the alternative ecotypes growing within each environment (local vs. foreign criterion) and for the effect of the environment on each ecotype (home vs. away criterion; Kawecki & Ebert, 2004). We used generalized linear mixed effect models (GLMMs) implementing the fitness component as response variable and ecotype, transplant environment, and their interaction as predictor variables, using a binomial error distribution for the categorical variables (i.e., flowering probability and survival probability) and zero-inflated Poisson distributions for seed count. We further analysed survival throughout the life cycle using mixed effect Cox models, which perform proportional hazards regression of time to event data with implementation of random effects. We fitted the Cox models with ecotype, transplant environment and their interaction as predictor variables using the package survival 2.44 (Therneau & Grambsch, 2000).

### Life-history variation

To elucidate whether adaptation is accompanied by life-history differences we estimated elasticities of individual vital rates and compared stable age distributions of each ecotype growing under contrasting environments. Elasticities estimate the proportional sensitivities of a change in a specific vital rate to population growth rate and their distribution through the life cycle describes the relative influence of vital rates on the population growth rate of each ecotype in a given environment. Stable age distributions describe the proportions of individuals in in each age class. Values and confidence intervals were inferred from the same matrix population models and bootstrapping procedures as described above.

### Impact of plant size on fitness

To investigate the impact of plant size on fitness, we implemented the fitness components at each life stage as a function of plant size at the previous stage, together with ecotype, transplant environment and their interactions as predictor variables in GLMMs. To gain an overall estimate of the effect of plant size across the life cycle, we standardized plant size and seed count within life stage and site to a mean of 0 and standard deviation of 1. We then used standardized plant size as predictor variable while considering each repeated measure of the same individual plant as a unique sample and included life stage as a covariate. For the modelling of the effect at subsequent life stages we used the non standardized data. We used binomial error distribution for the modelling of survival and flowering probability and a zero-inflated Poisson distribution for seed count. For the modelling of seed count we assumed that the probability of a structural zero varies with plant size (Brooks *et al.,* 2017). For the GLMMs and the Cox models, we included seed families nested within population and block nested within site as random effects, except in a few cases when low variation caused singular fits of the models (see results). We applied mixed effect models using the package lme4 v1.1 (Bates *et al.,* 2015) and zero-inflated models using glmmTMB v. 1.3 (Brooks *et al.,* 2017). We assessed significance levels of the interactions using likelihood ratio tests with package lmerTest v.3.3 (Kuznetsova *et al.,* 2017) and obtained estimates and significance levels of fixed effects with package emmeans v.1.5 (Lenth, 2017).

### Recombinant populations

A principal component analysis (PCA) based on the ddRAD data identified population structure in the F2 plants that were grouped them into three genetic clusters (see Pålsson, 2023a). In all statistical models of the F2 data described below, we accounted for variation between these clusters by including them as a covariate in the models and tested their interaction with other predictors, as well as their effect on the response variables. The genetic clusters were included in the final, most parsimonious, models if either the interaction with other predictors or the effect on the response variables were statistically significant as determined with likelihood ratio-tests (LRTs). Further, we accounted for the effect of transplant blocks as a random effect when this exerted a significant effect on the response variable. To improve model fit, for all studied traits, as well as the fitness proxies cumulative seed weight and seed count, values exceeding two standard deviations from the mean within each genetic cluster and site were excluded from the analyses (i.e., low environment cluster 1: 3, cluster 2: 7, cluster 3: 6; high environment, cluster 1: 6, cluster 2: 7, cluster 3: 2).

We aimed to compare selection estimates of plants at the same stage of the life cycle and therefore used data from the first year of flowering, i.e., 2019 and 2020, for plants growing in the low and high site, respectively. We investigated whether the phenotypic variation expressed in our F2 populations encompassed the phenotypic variation of the wild populations and if the dependence of flowering time and plant high on plant size was recapitulated in the F2 populations. We produced density plots of the trait distributions of the low and high elevation ecotypes and the F2s, growing at both elevations. For this visualization we used the mean flowering time, plant height and start of the season plant size across the experiment for the wild populations. We examined the genetic correlations between the traits by fitting plant height and flowering time as response variables and plant size, transplant site and their interaction as predictor variables in LMMs, using a Gaussian error distribution.

### Cumulative fitness estimates for phenotypic selection analyses

To obtain a cumulative estimate of relative fitness for the phenotypic selection analyses, we employed the aster modelling framework, which accounts for the hierarchal structure of fitness by incorporating multiple co-dependent components. Following Geyer et al., (2007), we modelled separate fitness components into hierarchal life-history stages following a graph structure (Figure S4). We used survival probability, flowering probability and seed production as the three layers of our models. Each layer contained two nodes representing performance in the individual fitness components during 2019 and 2020, where the performance in each of the nodes depends on the predecessor node. The binary variables survival and flowering probability were modelled with a Bernoulli error distribution and the seed count data with a Poisson error distribution. We scaled flowering time to start at 1 in both sites to account for the different start of the growing season in Julian days between the sites. We globally standardized flowering time, plant height and plant size to a mean of zero and standard deviation of one. We then identified the most parsimonious model within each site using a step-wise model selection approach by testing full models including all relevant two-way interactions against reduced models using LRTs. The full models included as predictor variables flowering time, plant height, plant size, genetic cluster, as well as two-way interactions between flowering time and plant height, and between each trait and genetic cluster. We subsequently used the output of the most parsimonious models to predict the expected fitness for each individual plant in each site separately. Aster models were fitted using the package Aster v1.1-1 (Geyer *et al.,* 2007). We relativized the cumulative fitness obtained from the aster models to the global mean across sites in line with recommendation for selection analyses that can be assumed to be non-frequency dependent (De Lisle and Svensson, 2017).

#### Phenotypic selection analyses

We estimated selection differentials, i.e., total selection acting on each trait, through cumulative fitness from the aster models by implementing it as response variable in LMMs. We modelled fitness functions using each trait separately as predictor variable, and included site and the interaction with site as covariate, using a Gaussian error distribution. Significance and effect sizes of the differentials were estimated from parametric bootstrap (n=5000) models and included either a linear or a quadratic predictor. The quadratic regression coefficients were subsequently doubled to obtain the quadratic selection differentials (Stinchcombe *et al.,* 2008).

To identify the targets of selection and dissect how selection acts through different fitness components, we estimated selection gradients using multivariable LMMs for cumulative fitness and total seed weight and GLMM for the probability to produce seeds. We relativized total seed weight to the global mean across sites. These models included all traits simultaneously, as well as the interaction between flowering time and plant height as predictor variables. For cumulative fitness and relative total seed weight we used Gaussian error distribution, and for the probability to produce seeds, binomial error distribution. We modelled the data of the two sites together and used a parametric bootstrap (n=5000) to obtain the selection coefficients within each site simultaneously. Statistical significance was determined according to 95% bias-corrected confidence intervals (i.e., non-overlapping zero) estimated on the same parametric bootstrap replicates that were used to obtain the selection coefficients.

## Results

### Phenotypic divergence

The elevational ecotypes displayed phenotypic divergence in plant height, flowering time and plant size throughout the life cycle, although the effect depended on the growing environment. Plants originating from low elevation produced significantly taller stalks, and flowered later than the high ecotype in both environments, albeit flowering time divergence was only statistically significant in the high environment (Figure 1a and b, Table S6, Table S7). Plants of the low elevation ecotype grew larger the high ecotype when growing in their home environment, but this difference was not expressed at high elevation (Figure 1a, Table S8). The divergence in all traits was accompanied by yearly variation, particularly in the statistical significance of the contrasts, whereas general patters remained consisted across the life cycle (Figure S5, Figure S6, Table S6, Table S7, Table S8).

**Figure 1.**
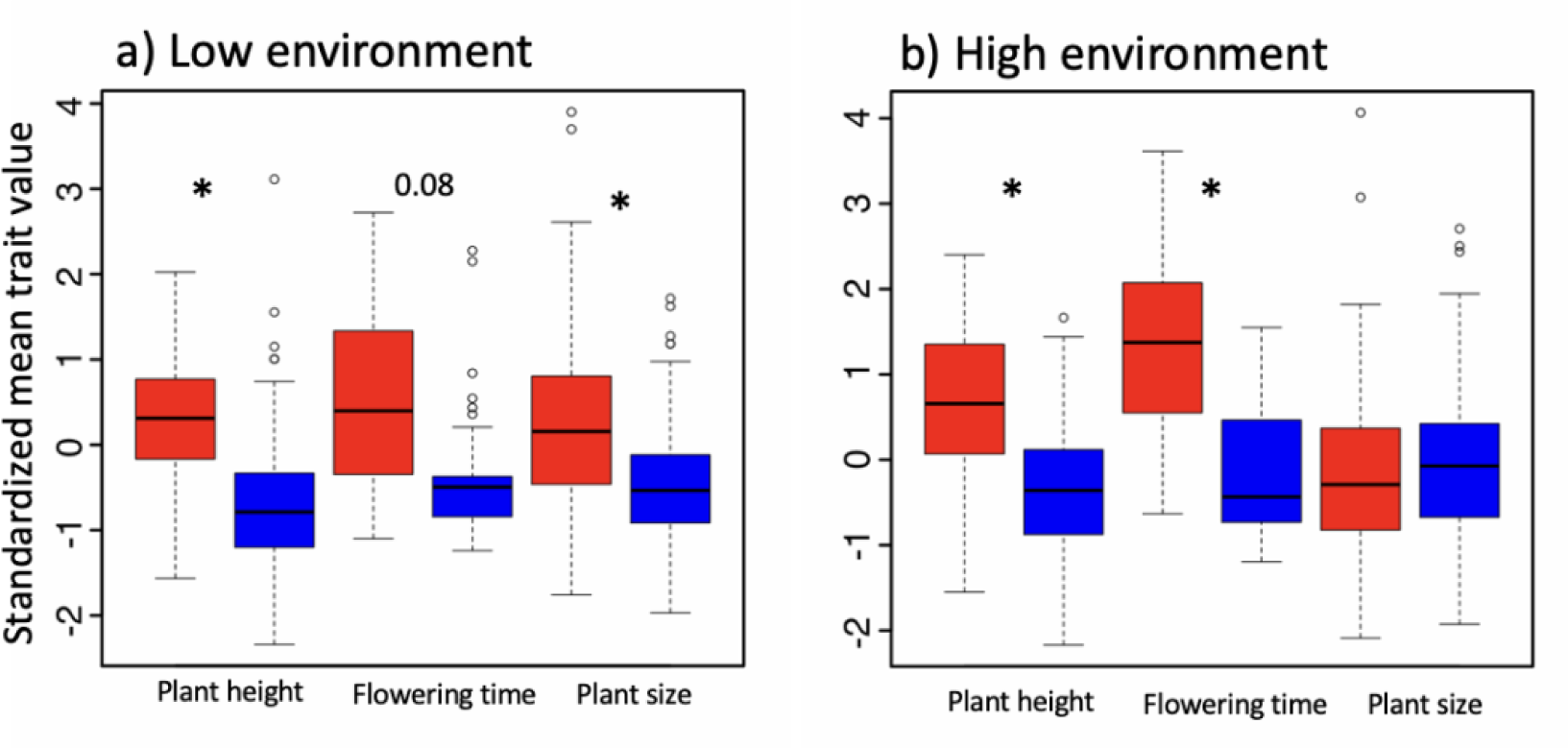
Phenotypic divergence in plant height, flowering time and plant size of the elevational ecotypes growing in the a) low and b) high environment. Boxes represent trait values standardized by life-stage and environment and statistical significance is inferred from linear mixed effect models. Red and blue denote the low and high elevation ecotypes, respectively. Significance of contrasts consistent to differential performance within each transplant environment are reported (**p<0.01, *p<0.05).

#### Genetic correlations between traits

For both ecotypes, genetic correlations underly a positive effect of plant size on plant height (Figure 2a – b, Table S9). Furthermore, larger plants of the low elevation ecotype flowered earlier in both environments (Figure 2c – d, Table S9). The traits dependence on plant size displayed strong yearly variation. Although not always statistically significant, the direction of the trends was consistent with the patterns observed in the mean values in all years except year five (Figure S7, Table S9).

**Figure 2.**
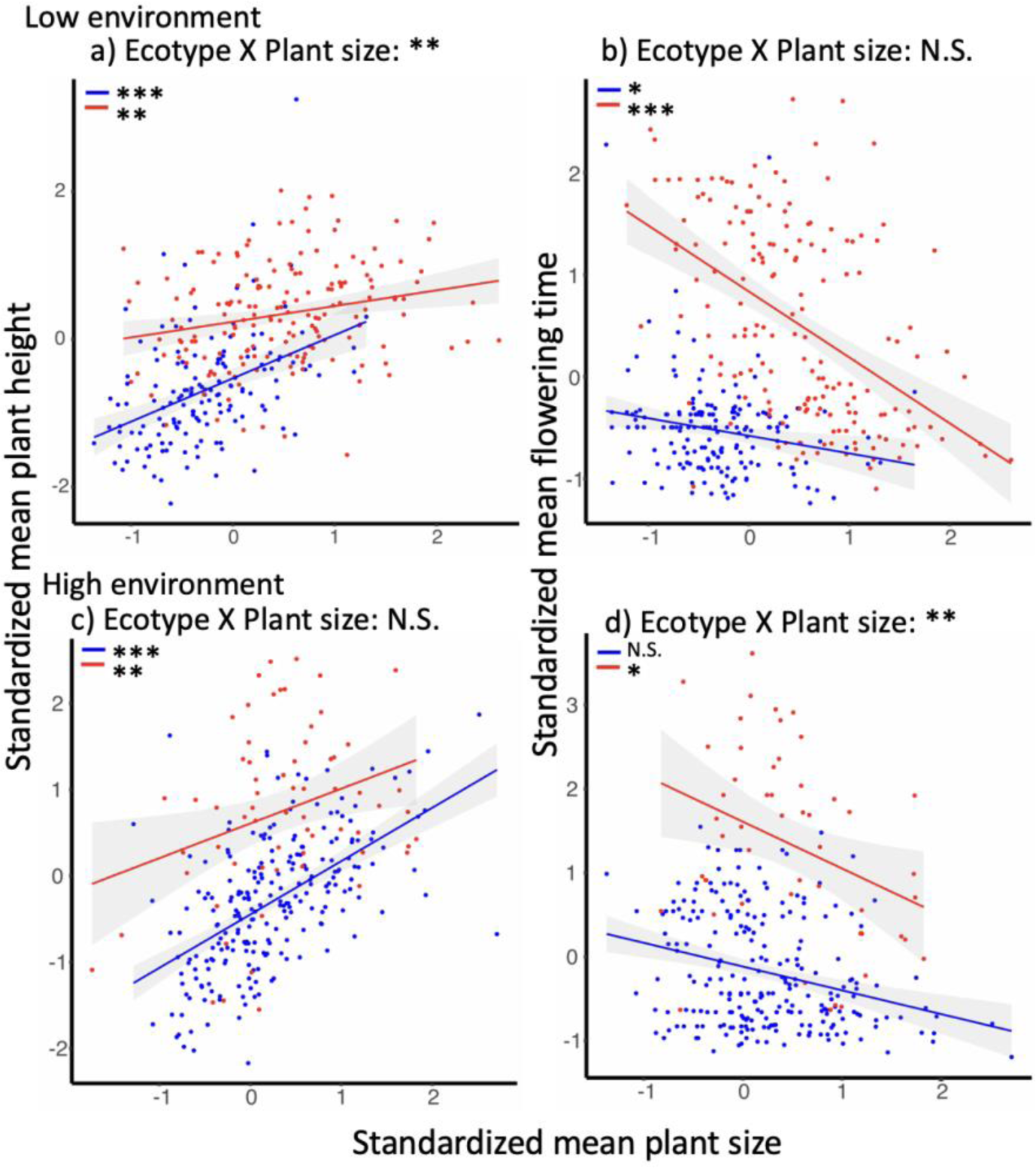
Relationship between plant height, flowering time and plant size of elevational ecotypes growing in the low and high environments. Left panels (a, c), relationship between plant height and plant size. Right panels (b, d), relationship between flowering time and plant size. Trait values are standardized by life-stage and environment and statistical significance is inferred from linear mixed effect models. Red and blue denote the low and high elevation ecotypes, respectively. Significance of the two-way interaction between ecotype and plant size and relationships between traits within each ecotype and transplant environment are reported. Short red and blue lines denote statistical significance of trait correlations for the low and high elevation ecotypes, respectively (***p<0.001, **p<0.01, *p<0.05).

### Elevational adaptation

Population growth rates estimated from MPMs revealed a significant advantage of the local over the foreign ecotype in both environments. The local ecotype always showed significantly higher population growth rate than the foreign (low environment: low ecotype: 1.5 (CI: 1.437, 1.575), high ecotype: 1.03 (CI: 0.968, 1.113); high environment: low ecotype: 0.815 (CI: 0.764, 0.870), high ecotype: 1.08 (CI: 1.042, 1.112)) (Figure 3a and b). The LTRE analyses revealed that the strongest negative impact of the vital rates of the foreign ecotype in low environment was reproduction the third year and fifth year whereas in the high environment, early survival and fourth and fifth year’s reproduction had a stronger relative impact (Figure 4d and e, Table S10).

**Figure 3.**
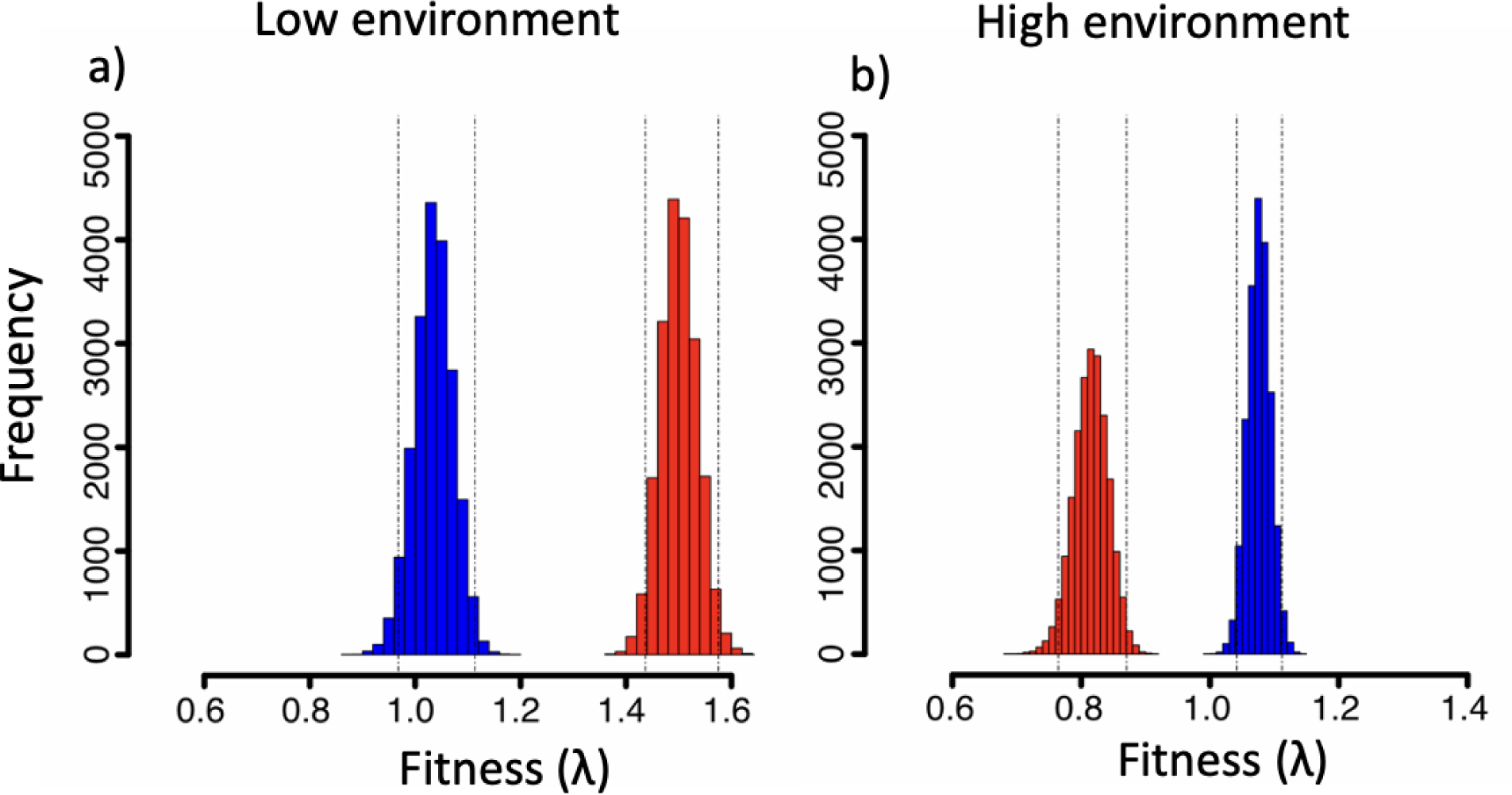
Integrative estimates of ecotype performance expressed as population growth rates (λ) growing in the transplant environments. a and b) Histograms representing population growth rate distributions based on 20 000 bootstrap replicates for the low (red) and high (blue) elevation ecotypes growing in the in the low and high environments. Dotted lines indicate bias corrected 95% confidence intervals.

**Figure 4.**
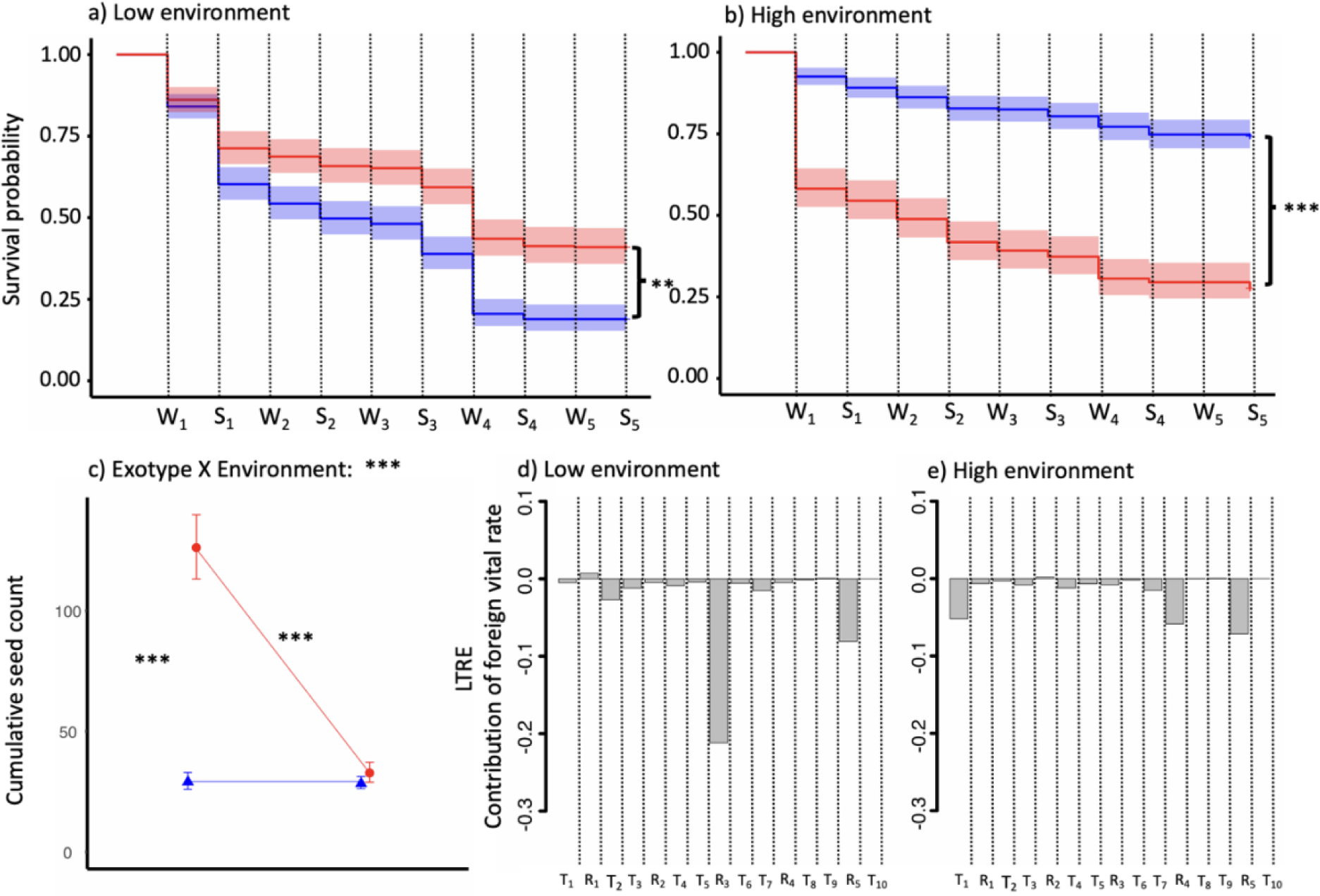
Contribution of individual fitness components to adaptation in the elevational environments. A and B) survival curves representing survival rates of the low (red) and high (blue) elevation ecotypes throughout the experiment at subsequent stages of the life cycle in the low a) and high b) environment. W_i_ and S_i_ denote the life stages (W_i_ winter survival and S_i_ summer survival). Asterisks at the side of each plot indicate significant cumulative differences between the survival curves of the two ecotypes as inferred from cox proportional hazard models and the shaded areas represent 95% confidence intervals. c) Performance in cumulative seed count of elevational ecotypes growing in the low and high environments. Symbols indicate mean estimate values inferred from generalized linear mixed effect models and bars indicate standard errors. Mean values are connected by reaction norms depicting the effect of the environment on each elevational ecotype. Red and blue colors denote the low and high ecotypes. Significance of ecotype by environment interactions and contrasts consistent to the local vs. foreign and home vs. away criteria are reported. d and e) LTRE showing the relative contribution of the vital rates expressed as survival (T_i_) and reproduction (R_i_) of the foreign ecotype to population growth rate at subsequent stages of the life cycle in the low d) and high e) environments. _i_ Indicate specific vital rates and life stages. (***p<0.001, **p<0.01).

Analyses of individual fitness component showed that survival has a strong impact on adaptation, as significant ecotype by environment interactions result in the local ecotype performing significantly better than the foreign ecotype in both environments (Figure 4a and b, Table S11). This difference was particularly pronounced in the high elevation environment, where the proportion of surviving local ecotypes was approximately double by the end of the experiment. Separate comparisons at subsequent stages of the life cycle identified significant ecotype by environment interactions, thus supporting these findings (Figure S8, Table S12).

Comparisons of cumulative seed count revealed a significant ecotype by environment interaction and strong signal for adaptation in the low elevation environment (Figure 4c, Table S13). In contrast, the two ecotypes yielded similarly low seed count at high elevation. The low ecotype produced significantly more seeds when growing in its home environment. We further detected strong inter seasonal variation in both seed count and flowering probability, with overall greater seed production in the third and fifth year (Figure S9, Figure S10, Table S13 and Table S14).

#### Elasticities and stable age distributions depend on ecotype and environment

The elasticity values extracted from the MPM showed overall strong influence of survival (i.e. T_i_) throughout the life cycle for both ecotypes in both environments (Figure 5a and b, Table S15). Elasticities of the reproductive vital rates were low for most years, with larger effects only in the third and fifth year in the low and high elevation environment, respectively. Estimates across ecotypes growing in the same environment were similar, with no statistically significant difference for any vital rate.

**Figure 5.**
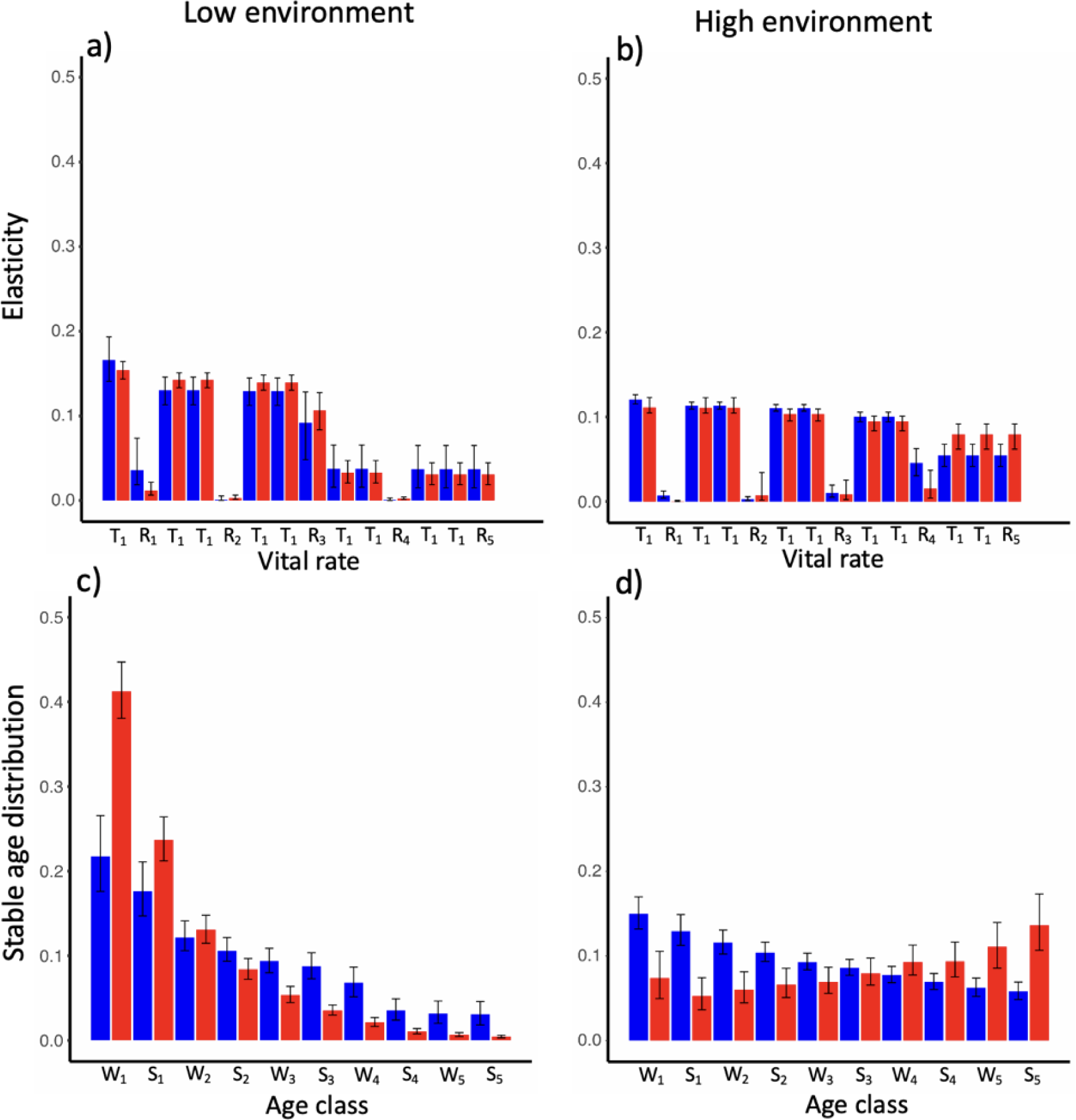
Environmental response of life-history traits expressed by the elevational ecotypes growing in the transplant experiment. a and b) Influence of the specific vital rates (i.e., elasticities) of survival (T_i_) and reproduction (R_i_) on population growth rates. c and d) Stable age distribution at subsequent summer (Si) and winter (W_i_) seasons. Elasticity and stable age distribution values are the mean of 20 0000 bootstrap replicates and error bars indicate bias corrected 95% confidence intervals. _i_ Indicate specific vital rates and life stages and red and blue bars indicate estimates for the low and high ecotype, respectively.

The stable age distributions revealed that in the low elevation environment, populations of both ecotypes, but in particular of the low ecotype, will consist primarily of young individuals (Figure 5c and d, Table S16). At high elevation we detected a divergent pattern between the ecotypes. The distributions of the low ecotype shifted towards an increased proportion of older individuals whereas the age distribution of the high ecotype remained skewed towards younger individuals.

#### Plant size affects performance in single fitness components

Plant size tended to have a positive effect on individual fitness components. In both environments, regardless of ecotype, larger plants were both more likely to flower and to survive (Figure 6a – d, Table S17, Table S18). Larger plants also produced more seeds, but in each environment, this effect was significant only in the local ecotype (Figure 6e and f, Table S19). Separate analyses at the subsequent life-stages revealed that impact of plant size on survival probability was statistically significant only for the time points when plants experienced strong mortality events (Figure S11, Figure S8). In these cases, the effect was consistently stronger for the foreign ecotype, except in the high environment the second winter (W2; Table S18). Plant size overall positively affected the probability of flowering during the first four years of the experiment, but not during the fifth year, and was consistently associated with increased seed output (Figure S12, Figure S13).

**Figure 6.**
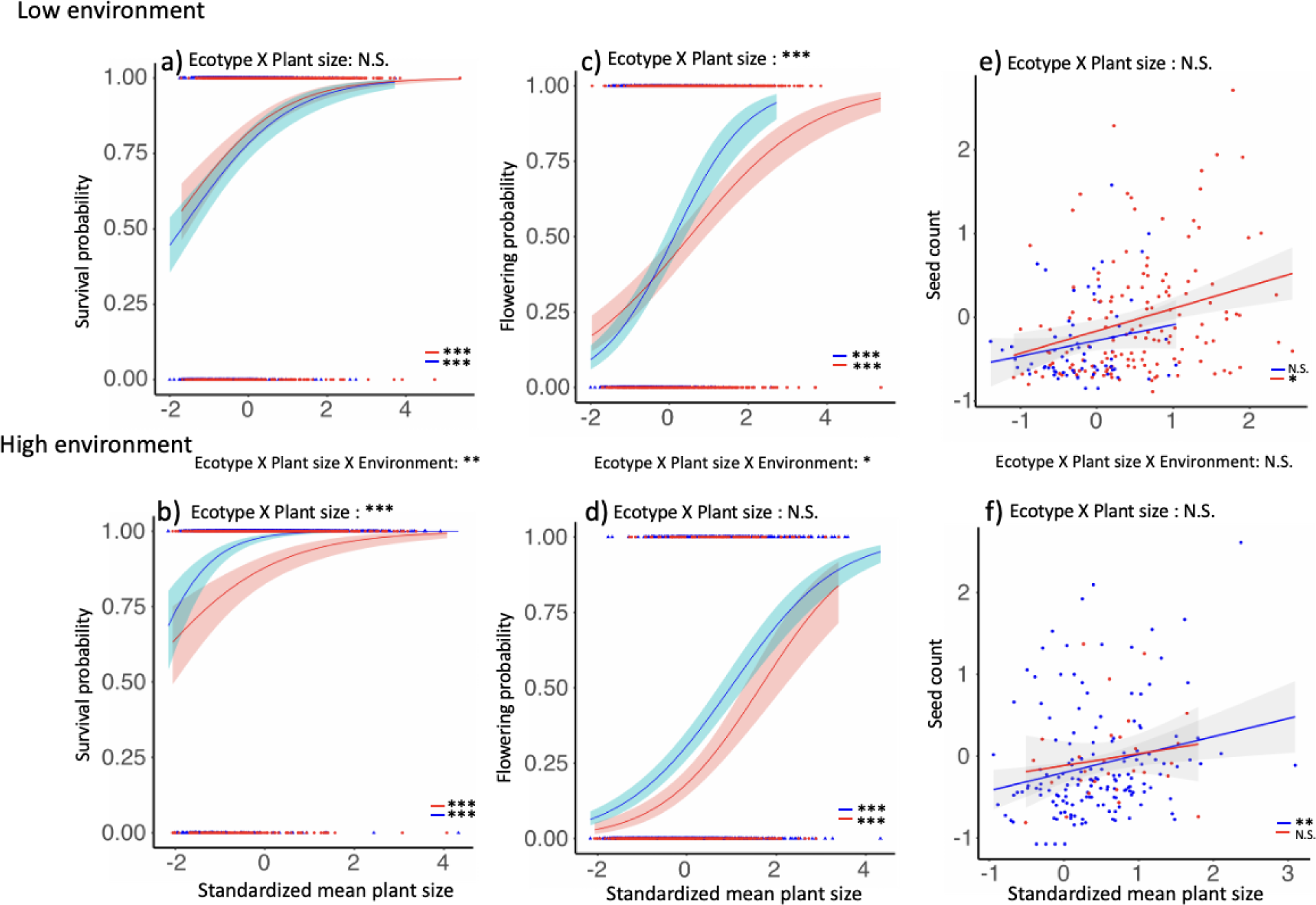
The effect of plant size on survival probability, flowering probability and seed count of elevational ecotypes growing in the low (a, c, e) and high (b, d, f) environment. Plant size and seed count are standardized by life-stage and environment. Red and blue lines indicate predicted relationships from generalized and linear mixed effect model regressions with 95% confidence intervals for the low and high elevation ecotypes, respectively. Corresponding red and blue triangles indicate values of plant size. Significance of the three-way interaction between ecotype, plant and the environment and of the ecotype X plant interactions and of the relationship between plant size and the separate fitness components are reported. Short red and blue lines denote statistical significance for the low and high elevation ecotypes, respectively (***p<0.001, **p<0.01, *p<0.05).

### Recombinant populations

The phenotypic distribution present in the wild populations was partly covered in the F2 populations (Figure S15). The dependence of plant height and flowering time on plant size identified in the wild populations was recapitulated in the F2 populations (Figure S14, Table S20).

We detected significant directional selection differentials acting towards earlier flowering and larger plants in both elevational sites, and taller stalks in the high site (Table 1). For flowering time, we further identified significant negative and positive nonlinear selection in the low and high site, respectively, and significant positive nonlinear selection for plant height in the high site. Selection for larger size was stronger in the low than in the high site (Table 1).

**Table 1.** Estimates of linear (S) and nonlinear (C) selection differentials for plant size, flowering time and plant height at the low and high elevation transplant sites. Selection differentials are estimated based on parametric bootstrap (n=5000) using univariable regression models and include 95% bias corrected confidence intervals. Significant results (i.e., estimates whose 95% confidence intervals do not overlap zero) are in bold.

| Traits | Directional selection |  | Nonlinear selection |  |
| --- | --- | --- | --- | --- |
|  | S cumulative fitness |  | C cumulative fitness |  |
|  | Low site | High site | Low site | High site |
| Plant size | **1.29 (0.86, 1.70)** | **0.28 (0.18, 0.383)** | -1.78 (-4.97, 0.75) | 0.01 (-0.21, 0.27) |
| Plant height | 0.11 (-0.01, 0.23) | **0.3 (0.23, 0.37)** | -0.22 (-0.40, 0.002) | **0.15 (0.06, 0.25)** |
| Flowering time | **-0.35 (-0.44, -0.26)** | **-0.27 (-0.34, -0.21)** | **-0.23 (-0.39, -0.08)** | **0.2 (0.12, 0.3)** |

The selection gradients based on cumulative fitness revealed strong directional selection for larger size in the low site and selection for earlier flowering in the high site (Table 2). We found significant selection gradients for earlier flowering and taller plants through the probability to produce seeds at the high elevation site. In the low elevation site, we did not find evidence for selection acting through this fitness proxy. Based on relative total seed weight we inferred significant selection for larger plant size at both elevations.

**Table 2.** Estimates of linear (b) selection gradients for plant size, flowering time and plant height at the low and high elevation transplant sites. Selection gradients are estimated based on parametric bootstrap (n=5000) using multivariable regression models and include 95% bias corrected confidence intervals. Results for cumulative fitness, relative total seed weight and probability to produce seeds, are reported. Significant results (i.e., estimates whose 95% confidence intervals do not overlap zero) are in bold.

| Traits | $\beta$ cumulative fitness | |
| --- | --- | --- |
|  | Low site | High site |
| Plant size | **1.00 (0.49, 1.47)** | 0.02 (-0.06, 0.1) |
| Plant height | 0.13 (-0.07, 0.33) | 0.11 (-0.04, 0.25) |
| Flowering time | 0.02 (-0.17, 0.21) | **-0.24 (-0.36, -0.14)** |
| $\beta$ relative total seed weight | | |
| Plant size | **1.26 (0.03, 2.54)** | **0.61 (0.14, 1.1)** |
| Plant height | 0.01 (-0.35, 0.36) | 0.50 (-0.04, 1.02) |
| Flowering time | -0.07 (-0.37, 0.22) | -0.11 (-0.76, 0.56) |
| $\beta$ probability to produce seeds | | |
| Plant size | 0.50 (-1.68, 2.73) | -0.42 (-1.38, 0.39) |
| Plant height | -0.27 (-0.85, 0.29) | **1.48 (0.91, 2.13)** |
| Flowering time | -0.47 (-0.98, 0.03) | **-0.8 (-1.34, -0.34)** |

## Discussion

Understanding how ecotype formation proceeds following range expansion into new habitats requires uncovering how adaptive traits contribute to the evolutionary response to novel selection regimes. We combined transplant experiments of wild populations and recombinant crosses to uncover the evolution of key traits in *D. sylvestris* following the recolonization of low elevation habitats in the central Alps after the Last Glacial Maximum. We show that populations inhabiting the extremes of elevational gradients evolved phenotypic divergence with a strong genetic basis, accompanied by adaptation to the contrasting selection regimes at high and low elevations. This process was driven by selection acting through alternate fitness components in the two environments, and was tightly linked to variation in life history traits. As a response to the warmer climate, the low elevation ecotype evolved a shorter life cycle characterized by high reproduction, in contrast to ancestral strategies apt to ensure self-maintenance under colder climates that constrains plant physiological processes. This process was mediated by the evolution of a plastic response to the higher energy environment at the warm end of the elevational gradient, thus enabling plants to achieve larger size in response to strong direct selection. Combined with selection for early flowering at the cold end of the gradient, these genetic responses underly the phenotypic divergence observed between the elevational ecotypes.

Strong evidence of elevational adaptation in *D. sylvestris* emerged from population growth models, where the local ecotype consistently outperformed the foreign ecotype in both the high and low elevation environment. While these models provide compelling evidence of local adaptation as inferred from integrative fitness estimates for these perennial plants, they also allow dissecting the contributions of separate life stages to fitness and help formulate hypotheses about how selection acts in different environments (Caswell, 1989; Caswell, 2001; Peterson *et al.,* 2016; Goebl *et al.,* 2022). In *D. sylvestris*, adaptation in the low elevation habitat was primarily driven by reproduction in the third year, whereas at high elevation, contributions of different stages of the life cycle were more uniformly distributed. Dissecting ecotype by environment interactions of complementary fitness components revealed that reproduction confers a strong advantage of the local ecotype at low elevation, concomitant with a skewed age distribution towards young individuals with high reproductive output. Contrarily, contrasts of reproductive output at high elevation yield similar estimates for both ecotypes. Here, plant performance was characterized by a strong adaptive signal in survival resulting in age classes similarly represented in the population, which indicates that population growth at high elevation relies on a conservative strategy. These inferences are in line with strategies of ensuring self-maintenance which commonly prevails under colder alpine conditions (von Arx *et al.,* 2006; Šťastná *et al.,* 2012; Laiolo & Obeso, 2017; Rosbakh & Poschlod, 2018). On the other hand, the low elevation habitat selected for a life history strategy that favors investment in early reproductive fitness at the cost of shorter life cycles. This response is consistent with patterns commonly recovered along elevational gradients (Hautier *et al.,* 2009; Kim & Donohue, 2011; Kim & Donohue, 2013; Laiolo & Obeso, 2017; Pérez *et al.,* 2020). A recent inference similar to ours was recovered by Peterson *et al.,* (2020) who used a common garden experiment to expose annual and perennial *Mimulus guttatus* populations to experimental drought conditions, resembling our low elevation environment, and revealed that selection favored allocation to reproduction under warmer and drier conditions. Warmer habitats typically allow for longer growing seasons, resulting in an overall higher energy input and resource availability that allow for increased reproductive output and competitive ability (Stearns, 1992; Laiolo & Obeso, 2017).

The shift to a faster life cycle at low elevation is associated with an increase in plant size expressed by the local compared to the foreign ecotype across all stages of the life cycle. In plants, size is a common indicator of resource accumulation driven by metabolic rates as defined by resource availability and physical parameters (Poorter *et al.,* 2011; Younginger *et al.,* 2017). Among these, temperature and length of the growing season play a key role (Körner, 2003; Davies, 2006; Körner, 2006; Körner, 2015). In *D. sylvestris*, we found that plant size was positively correlated with plant height and with earlier flowering. Plant size was also positively associated with survival and reproduction. Thus, size acts as a key determinant of both reproductive traits and fitness. This is a frequently reported physiological relationship (Bonser & Aarssen, 2009; Weiner *et al.,* 2009; Younginger *et al.,* 2017; Cheplick, 2020; Fournier *et al.,* 2020; Proulx, 2021), and that similar correlations between plant size and fitness components were recovered across ecotypes of *D. sylvestris* suggests that altered genetic correlations do not justify the observed shift in life-history traits. The ability of the low elevation ecotype to increase its size was only expressed when growing in its home environment while remaining latent under the contrasting environmental conditions at high elevation. This environmental dependence indicates that the faster life cycle achieved by the low ecotype constitutes an inducible strategy set off by the climate at low elevation, where higher resource acquisition allows faster growth. We deduce that selection in the low environment acted on the genetic basis of the plastic response to the warmer climate, fueling a shorter cycle with resource acquisition devoted to intense reproduction.

At both elevations, total selection indicated that plants that flowered earlier, with tall stalks and large size had higher fitness, with particularly high coefficients for selection on size at low elevation. Multivariable analyses corroborate that this signal in plant size is the result of direct selection, while genetic correlations underlie the selection differentials estimated for the other traits. Notably, dissecting the impact of selection on separate fitness components revealed that direct selection for large size was driven by reproductive output, while the coefficient estimated for the probability to reproduce was close to zero. This is in line with the adaptive role of size at low elevation as a key driver of a fast life cycle where reproduction can be achieved by most individuals, but differential fitness is primarily driven by the investment in large reproductive output.

In the high elevation habitat, a contrasting scenario was recovered, as direct selection favored early flowering, while total selection estimated for plant size results from the strong genetic correlation between these two traits. Moreover, significant gradients were driven by the effect of selection on the probability to produce seeds rather than on reproductive output. Flowering time is a crucial transition that, for entomophilous plants like *D. sylvestris*, must coincide with the availability of pollinators and is necessarily linked to abiotic conditions allowing the reproductive stage of the life cycle (Elzinga *et al.,* 2007; Amasino, 2010; Ehrlén, 2015; Gaudinier & Blackman, 2019; Ehrlén *et al.,* 2020). Early flowering ensures successful reproduction in alpine environments where the short summer season constrains fruit maturity but can come at the cost of diminished resources that can be allocated to reproductive output (Obeso, 2002; Stinson, 2004; Gimenez-Benavides *et al.,* 2007; Anderson *et al.,* 2012; Hamann *et al.,* 2021). This trade-off can be exacerbated in low resource environments, or alternatively be reduced following the release from limitations imposed by the abiotic environment (Tuomi *et al.,* 1983; Obeso, 2002; Shefferson *et al.,* 2003; Sletvold & Ågren, 2015; Willi & Van Buskirk, 2022).

Our analyses of total selection acting on flowering time and plant size indicated that adaptation to low elevation evolved from relaxed trade-offs between the probability to produce seeds and reproductive output as mediated by the warmer climate. Importantly, selection gradients for large size at low elevation and early flowering at high elevation are in line with genetic divergence in phenotypic traits between ecotypes, which thus results from alternate evolutionary response to selection regimes at the extreme of the elevational gradient. On the other hand, despite our expectations informed by general patterns across plant systems (Halbritter *et al.,* 2018), we did not find evidence for direct selection acting on plant height. Selection for tall stalks is typically related to competition for light and pollinators, and is frequently observed in highly competitive environments (Gervasi & Schiestl, 2017; Zu & Schiestl, 2017; Halbritter *et al.,* 2018). However, *D. sylvestris* naturally inhabits rocky outcrops which are largely void of density dependent competition from direct neighbors, and long stalks are typically arcuate, thus favoring visits through lateral approach by pollinators. Thus, we infer that in contrast to plant size and flowering time, ecotypic divergence in plant height in *D. sylvestris* is not mediated by spatially divergent natural selection but is instead a consequence of genetic correlations with these adaptive traits.

The adaptive shift towards a shorter life cycle under warmer conditions in *D. sylvestris* recapitulates common trends observed across many plant species along temperature gradients. Our results shed light on processes that governed the recent response to climate driven selection in this species, and while mechanisms remain specific to each system, similar principles may underlie evolutionary trajectories shared across species. Our results show that the recent response of *D. sylvestris* to postglacial warming proceeded through selection acting on adaptive variants that underlie the plastic response of physiological processes associated with increased plant growth. These variants were likely present as standing genetic variation in refugial populations during the LGM and can be expected to be maintained in the gene pool of current high elevation populations descendant from recolonizing lineages. As anthropogenic climate change imposes novel selection regimes, this adaptive potential can form the basis of a continued evolutionary response through the realization of past evolutionary trajectories such as those described in this study. On the other hand, whether low elevation populations harbor the genetic potential for further evolution, and which trajectories this may imply, remains unknown. Our study is a first step to unravel past evolutionary responses to climate warming and further experiments are needed to uncover the existence of latent phenotypes that may be expressed and become adaptive under future climatic conditions.

## Acknowledgements

We would like to thank Ursina Walther for her contributions in the field experiments and valuable input on the statistical analyses, and Michael Gehrig for conducting the sowing experiment. We are grateful to Maja Frei, Esther Zürcher and Malwine Peter for their support with plant cultivation in the greenhouse. We are thankful to Pfyn-Finges natural park, Köbi Graven and Stefan Hardegger for allowing us to use their land for the transplant sites, and to members of the Plant Ecological Genetics group for their help with transplants in the field. This project was supported by the grants 31003A_160123 and 31003A_182675 from the Swiss National Science Foundation (SNSF) to AW.

## Author contributions

AP, SF and AW designed and set up the transplant experiments. AP collected the field data and performed all analyses. AP wrote the manuscript, which all authors revised.

