## Supplementary materials for "Trait evolution linked to climatic shifts contributes to adaptive divergence in an alpine carnation (*Dianthus sylvestris*)"

**Supplementary materials
Supplementary figures**

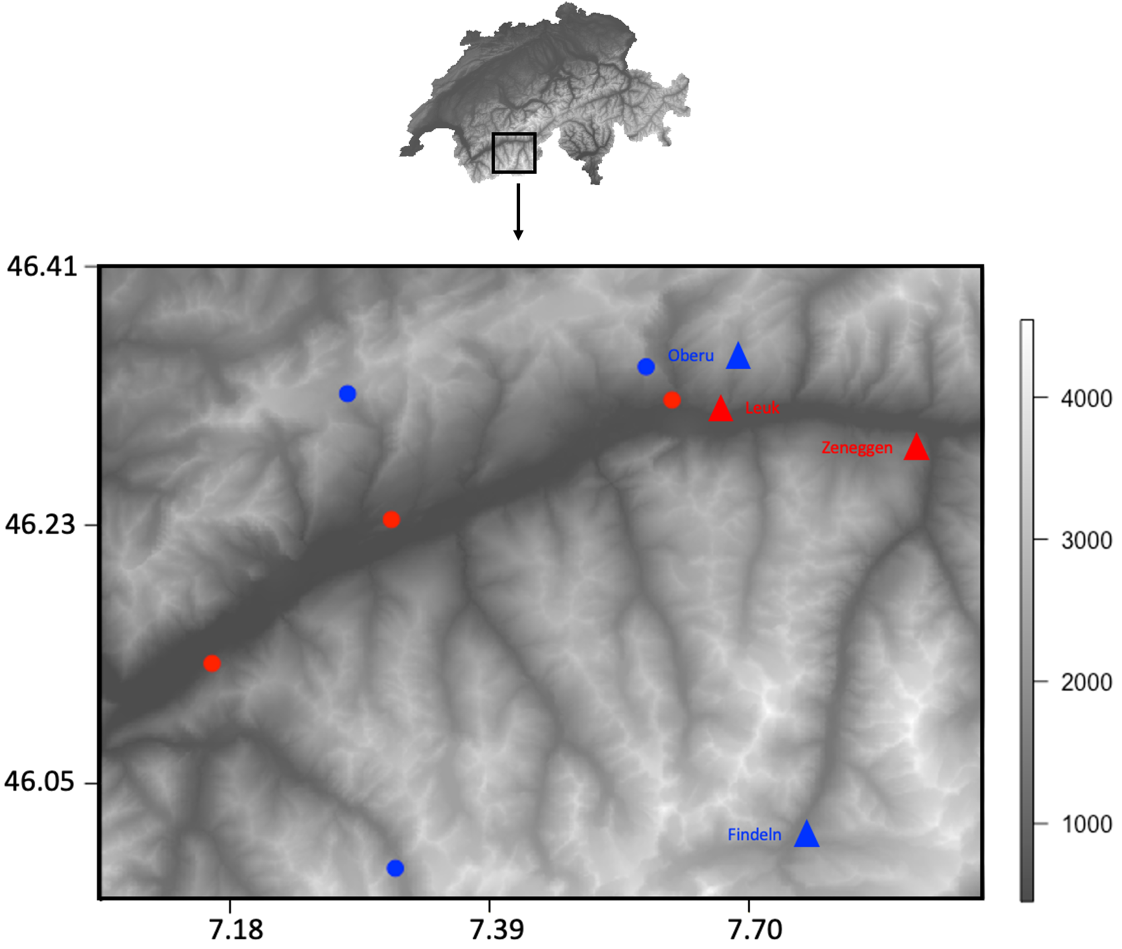

**Figure S1.** Map of the study area in the central Swiss Alps (Upper Rhône Valley). Locations of the three high (blue) and low (red) elevation populations of *D. sylvestris* (circles) and four transplant sites (triangles).

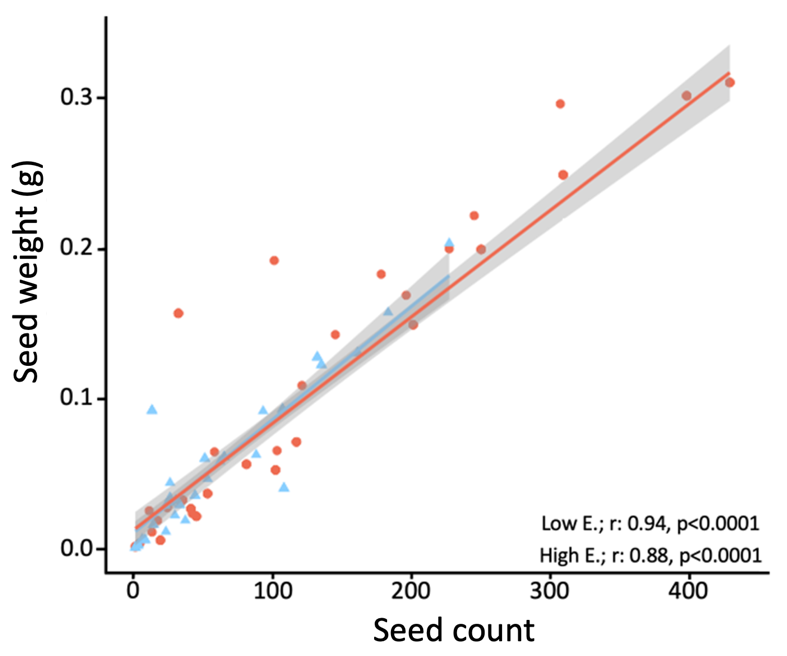

**Figure S2**. Correlation between seed weight (g) and seed count for seeds from 84 and 82 F2 individuals growing in the low (red) and the high (blue) elevation transplant site, respectively. Blue and red lines with 95% confidence intervals indicate the predicted relationship based on linear model regressions. Correlation coefficients (r) and p-values are reported.

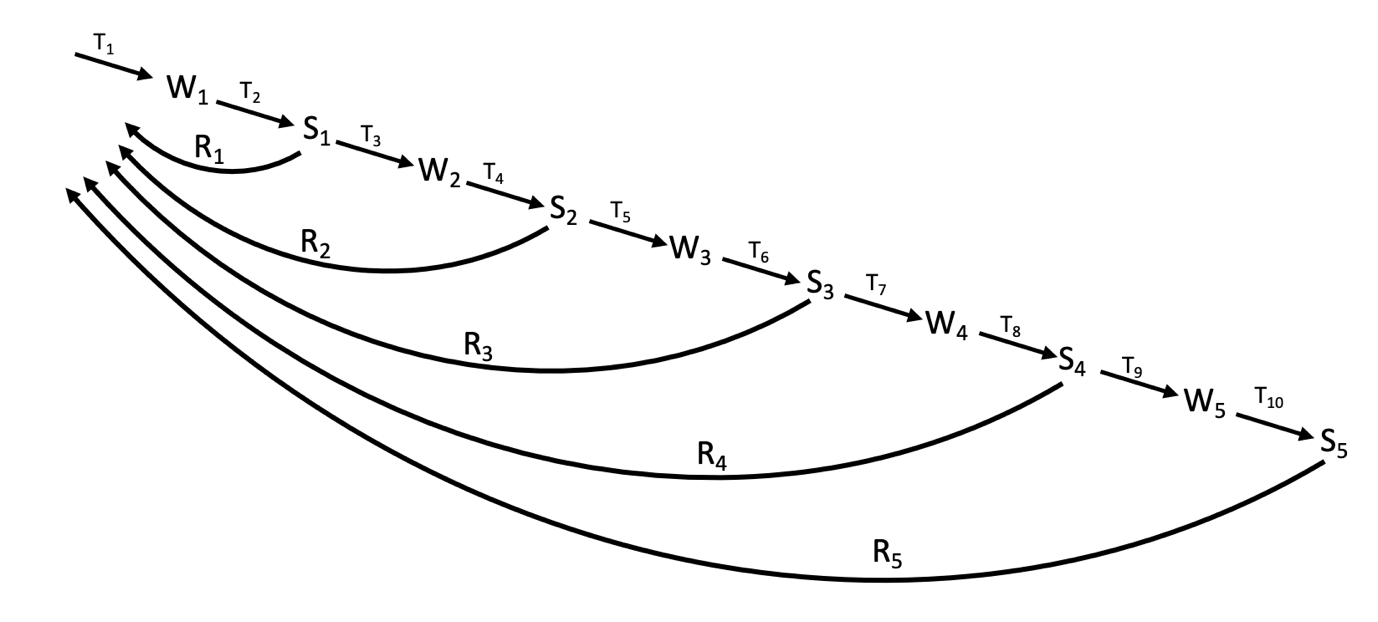

**Figure S3.** Age-classified life-cycle graph for *D. sylvestris* used for the matrix population models. The life cycle is divided into subsequent summer (S_i_) and winter (W_i_) stages, represented by indexes. The vital rates (i.e., survival T_i_ and reproduction R_i_) are inferred as transitions between the stages.

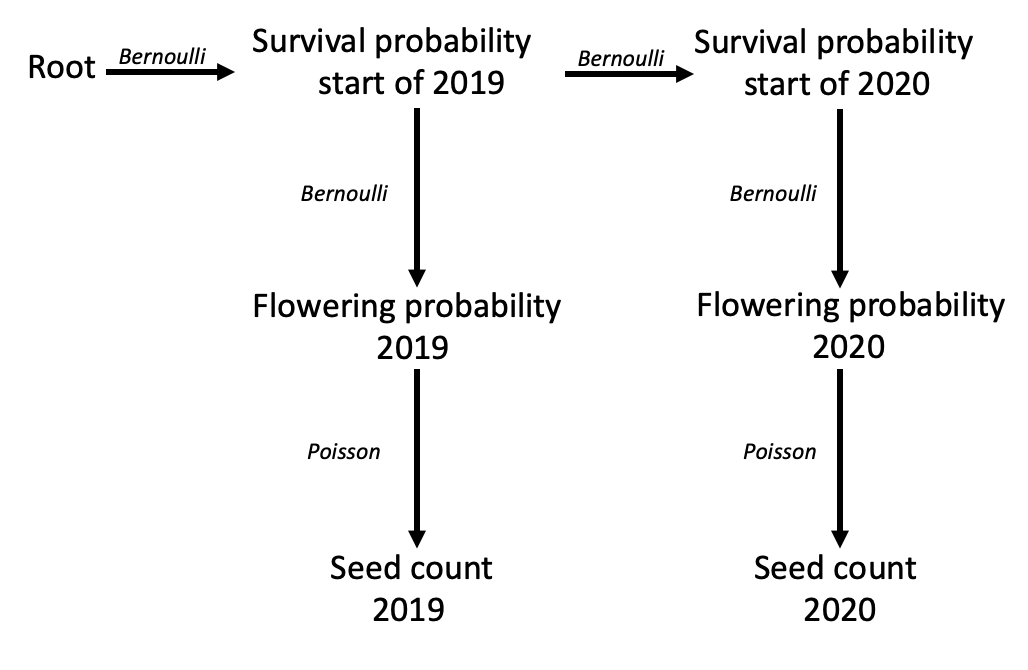

**Figure S4.** Life-history graph depicting the structure of the aster models. The graph consists of three layers representing, survival probability, flowering probability and seed count. Each node in the graph represents performance in the separate fitness components for the different seasons. The binary variables of survival and flowering probability were modelled using Bernoulli error distribution and the seed count data using Poisson error distribution.

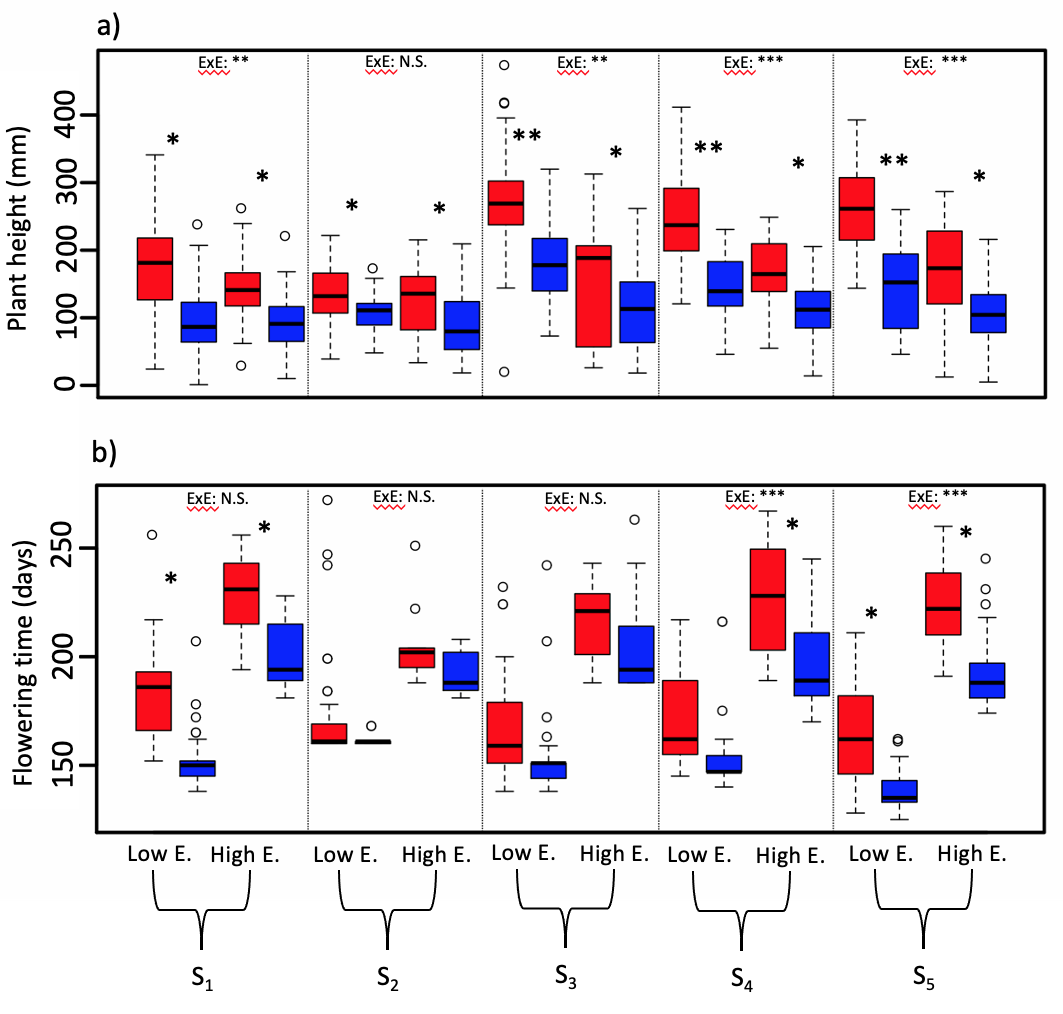

**Figure S5**. Phenotypic divergence in a) plant height (mm) and b) flowering time (days) of elevational ecotypes growing in the low and high environment at subsequent growing seasons. Boxes represent the raw values and statistical significance is inferred from linear mixed effect models. S_i_ denote the growing seasons. Low env. and High env. denote low and high environments, respectively. Red and blue denote the low and high elevation ecotypes, respectively. Significance of ecotype by environment interactions (ExE) and contrasts consistent to differential performance within each transplant environment are reported (***p<0.001, **p<0.01, *p<0.05).

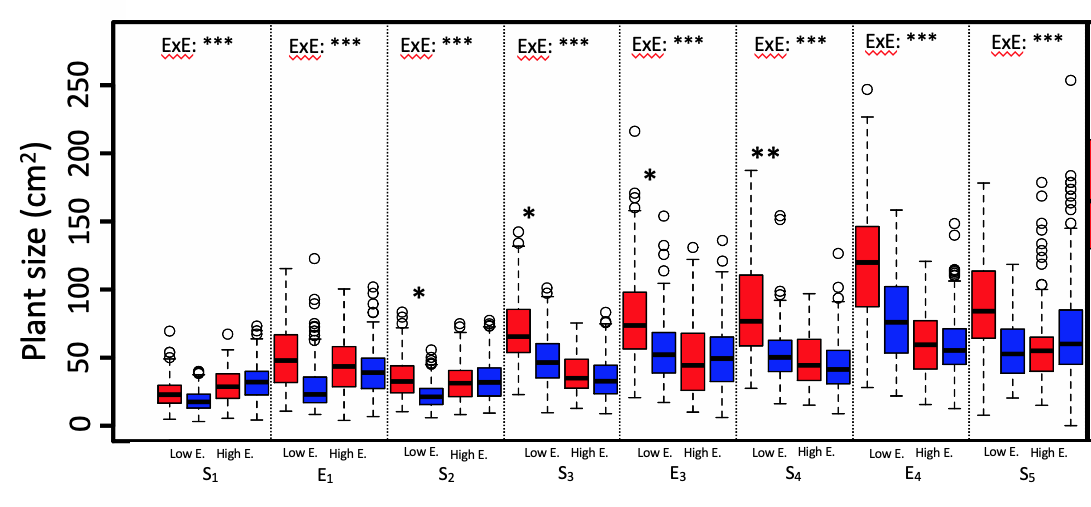

**Figure S6**. Phenotypic divergence in plant size (cm^2^) of elevational ecotypes growing in the low and high environment at subsequent stages of the life cycle. Boxes represent the raw values and statistical significance is inferred from linear mixed effect models. Low env. and High env. denote low and high environment, respectively. Red and blue denote the low and high elevation ecotypes, respectively. Significance of ecotype by environment interactions (ExE) and contrasts consistent to differential performance within each transplant environment are reported (***p<0.001, **p<0.01, *p<0.05).

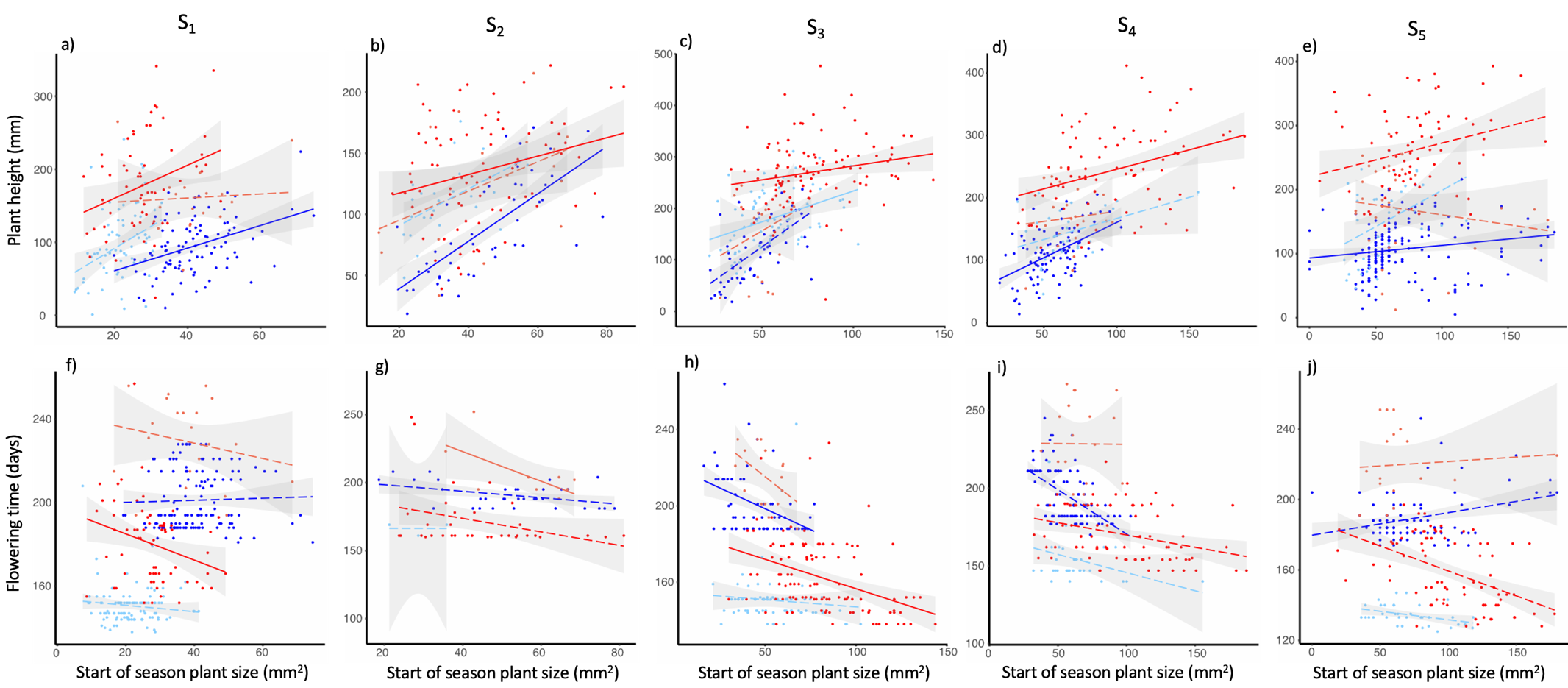

**Figure S7**. Relationship between plant height (mm), flowering time (days) and plant size (mm^2^) of elevational ecotypes growing in the low and high environments at subsequent growing seasons. Top panels (a-e), relationship between plant height and plant size. Bottom panels (f-j), relationship between flowering time and plant size. S_i_ denote the growing seasons. Dark red and blue indicate low and high elevational ecotypes growing in their home low and high environment, respectively. Light red and blue indicate low and high and high elevational ecotypes growing in their respective away environments. Filled and dashed lines indicate statistically significant (0.05>p) and nonsignificant relationships, respectively.

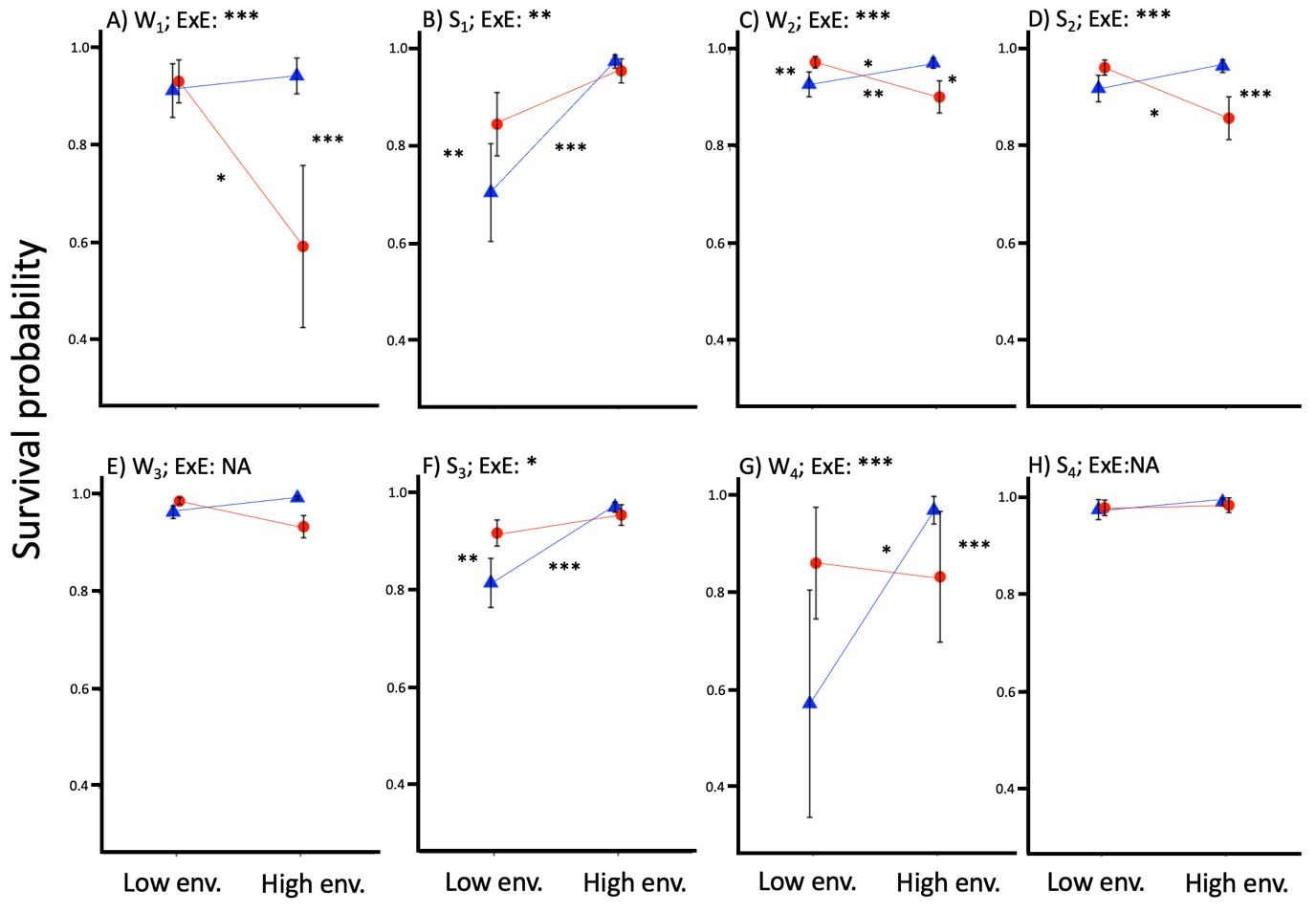

**Figure S8**. Performance in survival probability of elevational ecotypes growing in the low and high environment at subsequent growing seasons. Symbols indicate mean estimate values inferred from generalized linear mixed effect models and bars indicate standard errors. Mean values are connected by reaction norms depicting the effect of the environment on each elevational ecotype. Red and blue colors denote the low and high ecotype, respectively and Low and High env. indicate the low and high environments and. W_i_ and S_i_ denote the life stages (W winter survival and S summer survival). Significance of ecotype by environment interactions (ExE) and contrasts consistent to the local vs. foreign and home vs. away criteria are reported (***p<0.001, **p<0.01, *p<0.05).

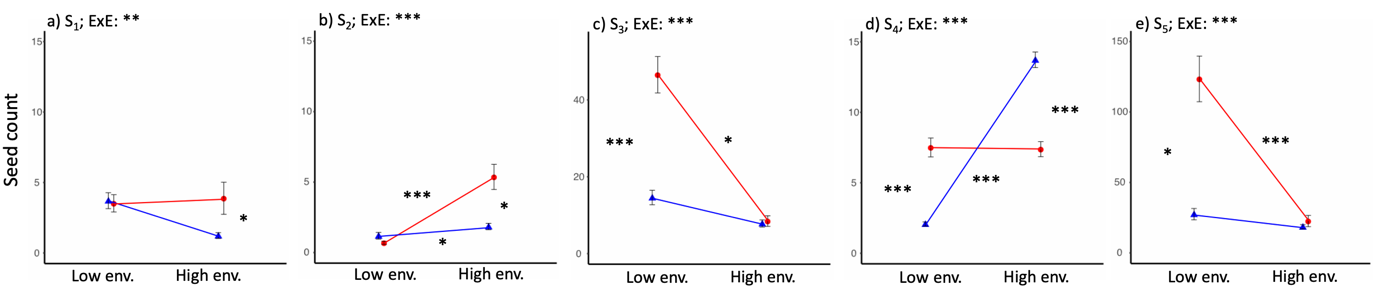

**Figure S9.** Performance in seed count of elevational ecotypes growing in the low and high environment at subsequent growing seasons. Symbols indicate mean estimate values inferred from generalized linear mixed effect models and bars indicate standard errors. Mean values are connected by reaction norms depicting the effect of the environment on each elevational ecotype. Red and blue colors denote the low and high ecotype, respectively and Low and High env. indicate the low and high environments and S_i_ denote the growing seasons. Significance of ecotype by environment interactions (ExE) and contrasts consistent with the local vs. foreign and home vs. away criteria are reported (***p<0.001, **p<0.01, *p<0.05).

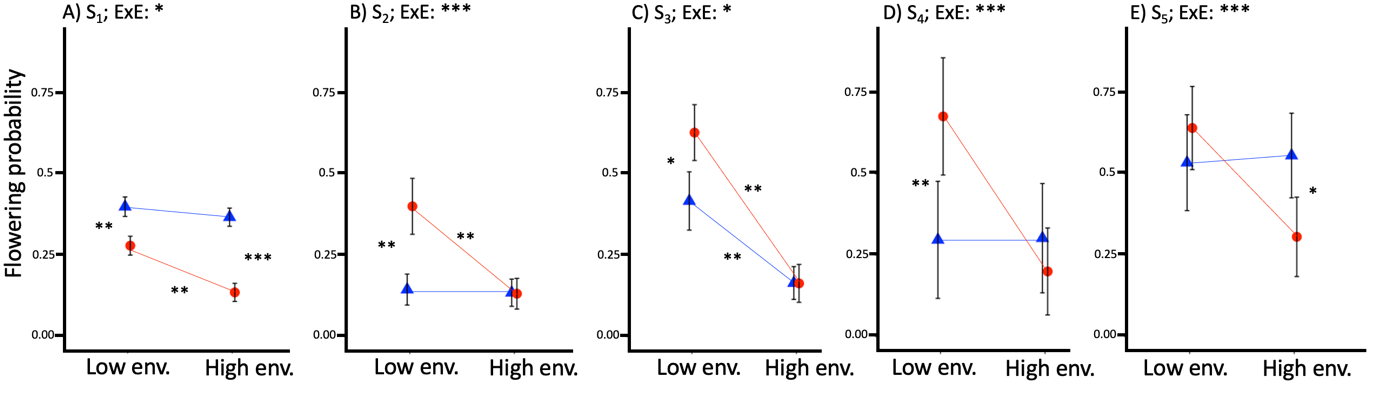

**Figure S10**. Performance in flowering probability of elevational ecotypes growing in the low and high environment at subsequent growing seasons. Symbols indicate mean estimate values inferred from generalized linear mixed effect models and bars indicate standard errors. Mean values are connected by reaction norms depicting the effect of the environment on each elevational ecotype. Red and blue colors denote the low and high ecotype, respectively, and low and high env. indicate the low and high environments and S_i_ denote the growing seasons. Significance of ecotype by environment interactions (ExE) and contrasts consistent with the local vs. foreign and home vs. away criteria are reported (***p<0.001, **p<0.01, *p<0.05).

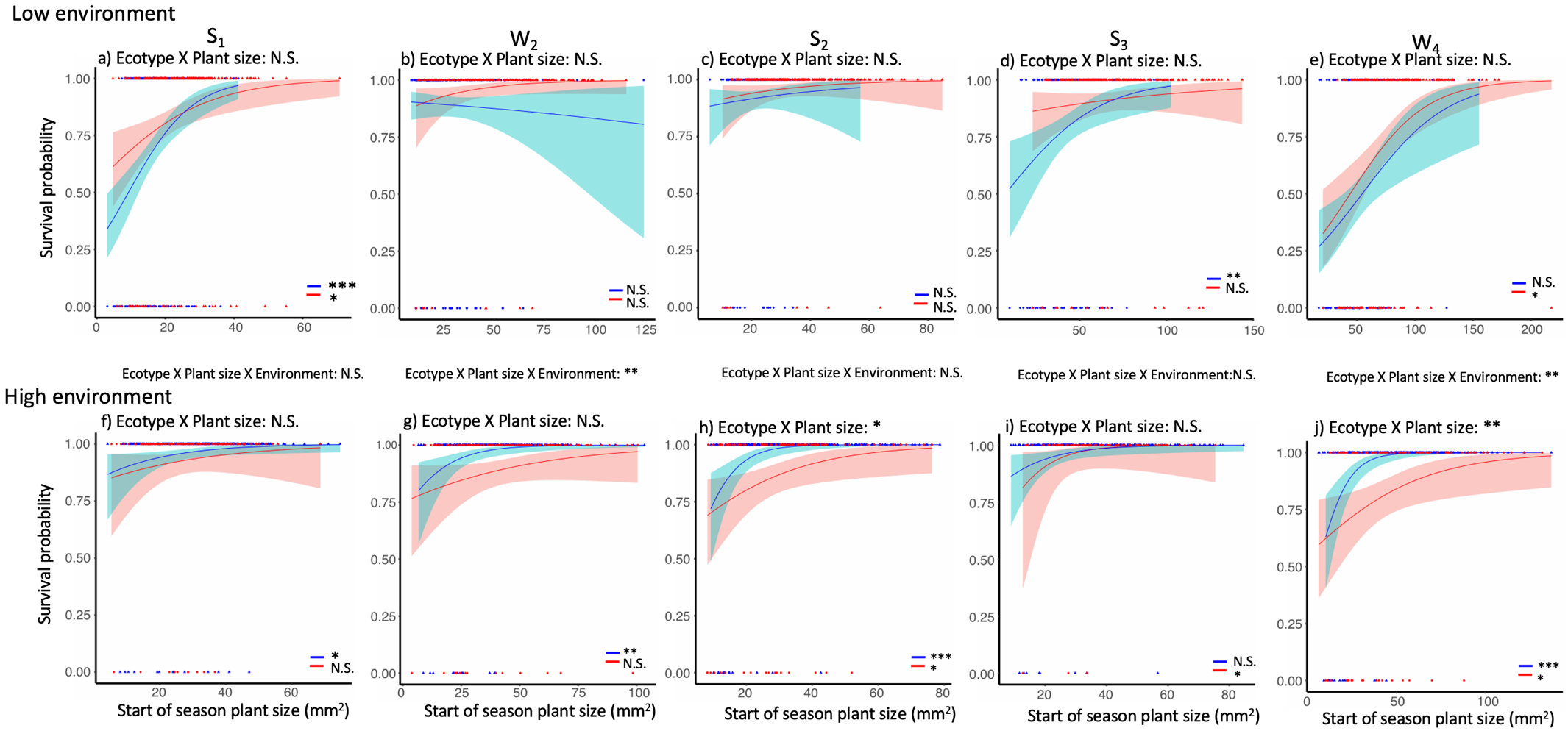

**Figure S11.** Effect of plant size on survival probability of elevational ecotypes growing in the low (a-e) and high (f-j) environments. W_i_ denotes the winter seasons. Red and blue lines indicate predicted relationships from generalized linear model regressions with 95% confidence intervals for the low and high elevation ecotypes, respectively. Corresponding red and blue triangles indicate empirical values of plant size at the start of each growing season. Significance of the three-way interaction between ecotype, environment and trait and ecotype by trait interactions within each environment and of the relationships between plant size at the start of the previous growing season and survival probability are reported (***p<0.001, **p<0.01, *p<0.05).

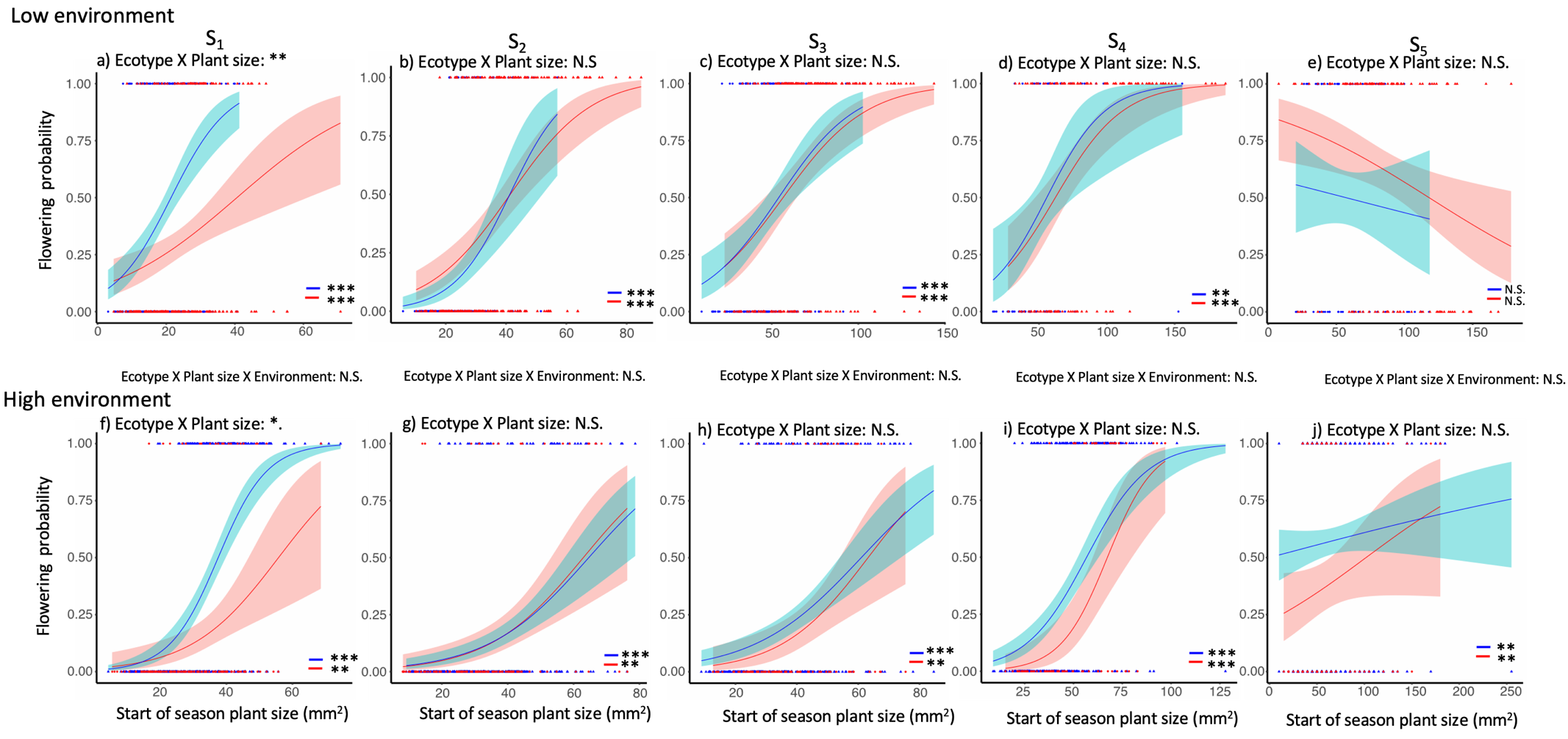

**Figure S12**. Effect of plant size on flowering probability of elevational ecotypes growing in the low (a-e) and high (f-j) environment. S_i_ denote the growing seasons. Red and blue lines indicate predicted relationships from generalized linear model regressions with 95% confidence intervals for the low and high elevation ecotypes, respectively. Corresponding red and blue triangles indicate empirical values of plant size at the start of each growing season. Significance of the three-way interaction between ecotype, environment and trait and ecotype by trait interactions within each environment and of the relationships between plant size at the start of each growing season and flowering probability are reported (***p<0.001, **p<0.01).

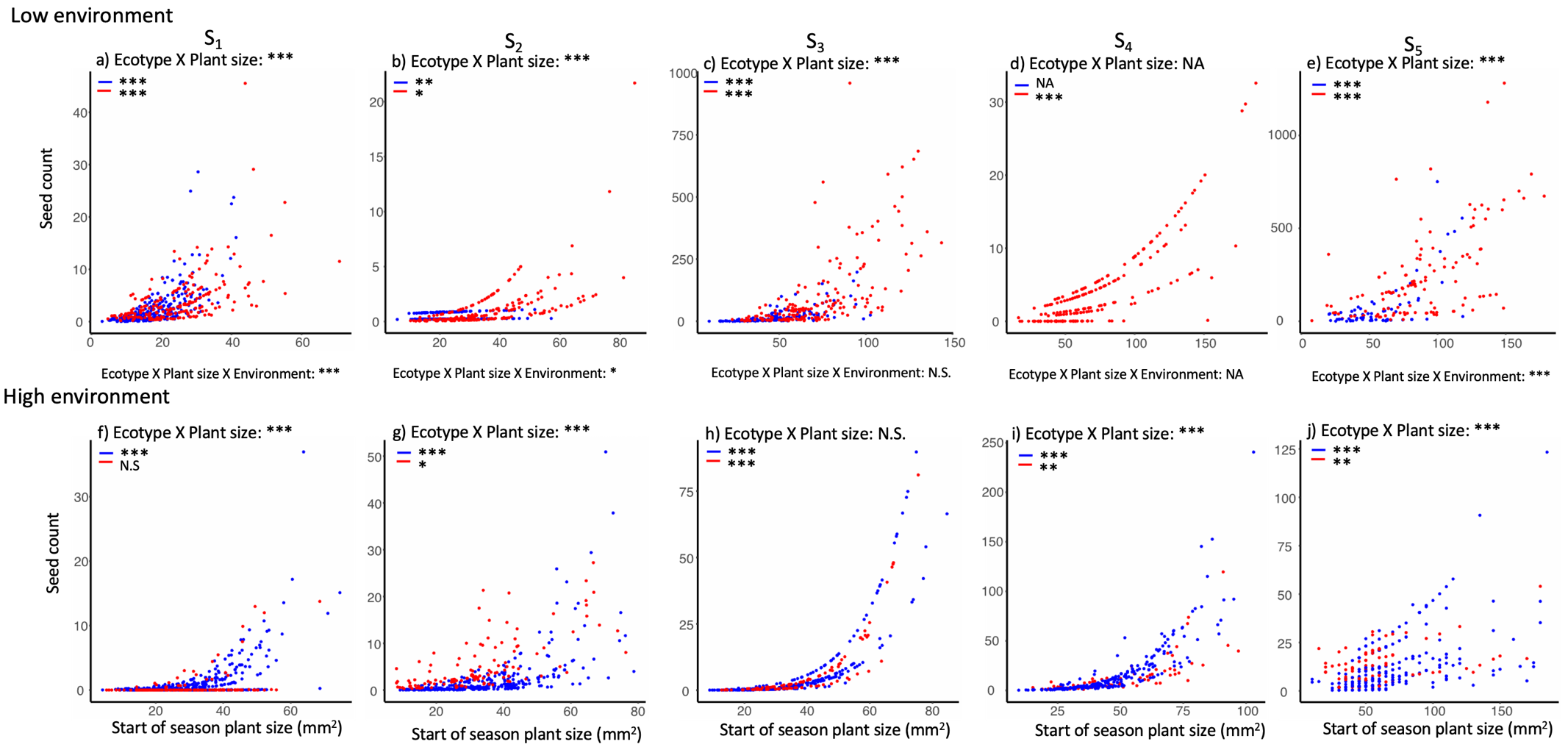

**Figure S13**. Effect of plant size on seed count of elevational ecotypes growing in the low (a-e) and high (f-j) environment. W_i_ denotes the winter seasons. Red and blue lines are added linear regression lines (with 95% confidence intervals), for the low and high ecotypes, respectively, on top of the relationship between plant size and seed count, estimated using zero-inflated poisson models. Significance of the three-way interaction between ecotype, environment and trait and ecotype by trait interactions within each environment and of the relationships between plant size at the start of the previous growing season and seed count are reported (***p<0.001, **p<0.01, *p<0.05).

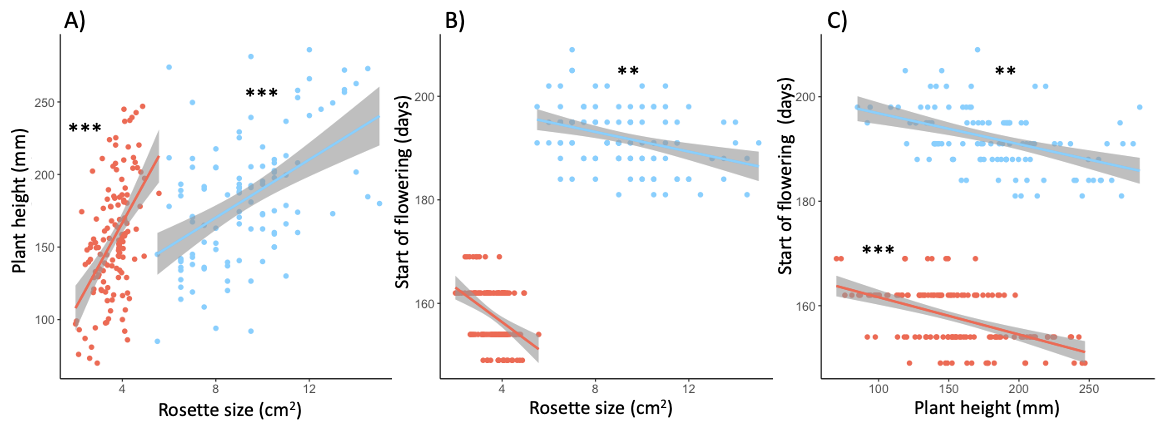

**Figure S14**. Relationship between plant size, plant height and flowering time at the low and high elevation transplant sites in 2019 and 2020, respectively. A) Plant height (mm) and plant size (cm^2^), B) flowering time (days) and plant size (cm^2^), C) flowering time (days) and plant height (cm^2^), Red and blue denote data from the low site in 2019 and the high site in 2020, respectively. Significance of the effect of the predictor trait (X axes) on the response trait (Y axes) as extracted from linear mixed effect models are reported (***p<0.001, **p<0.01)

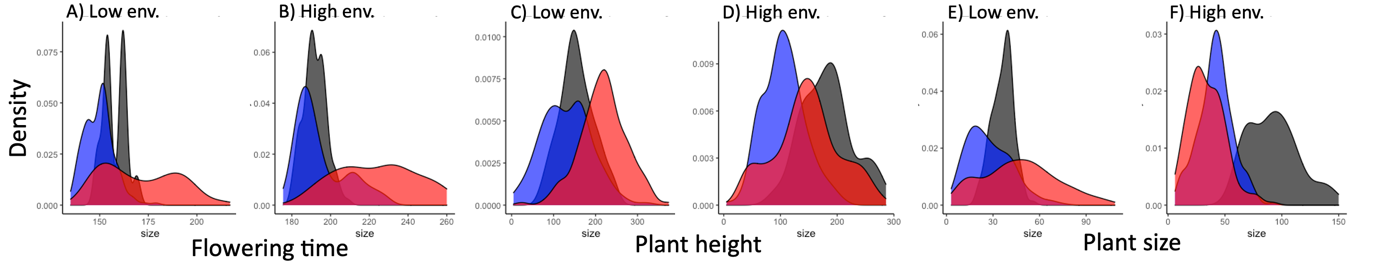

**Figure** **S15**. Density plots over trait distributions of wild and F2 populations growing in low and high environments and transplant sites (Low env. and High env.). Red, blue and black denote data of the low ecotype, high ecotype and the F2 populations, respectively.

**Supplementary tables**

| **Population** | **Elevation class** | **Elevation (m.a.s.l.)** | **Coordinates** | **Year sampled** |
| --- | --- | --- | --- | --- |
| Varen | Low | 768 | 46.32, 7.62 | 2012 |
| Saxon | Low | 570 | 46.13, 7.16 | 2012 |
| Saviese | Low | 746 | 46.24, 7.34 | 2014 |
| Chäller | High | 2062 | 46.34, 7.60 | 2012 |
| Tsanfleuron | High | 2110 | 46.32, 7.30 | 2014 |
| Val de Bagne | High | 2281 | 45.99, 7.34 | 2014 |

**Table S1.** Locations of the wild *D. sylvestris* populations included in the study. Population name, elevation class (low versus high), elevation (meters above sea level), coordinate (latitude, longitude) and year sampled are reported.

**Table S2.** Overview of the reciprocal transplant experiment of wild *D. sylvestris* populations. Transplant site name, elevational environment denominator (i.e., low or high), elevation (meters above sea level), coordinates (latitude, longitude), nr. of plants per population and transplant site and nr. of maternal families and mean nr. of plants per family (± SD) are reported. Note that all the plant nrs. refer to plants alive after transplant shock and hence form the basis of our analyses.

| **Site** | **Elevational environment** | **Meters above sea level** | **Coordinates: latitude, longitude** | **Population** | **Number of plants alive after transplant shock** | **Number of maternal families** | **Mean nr. of plants per family ± SD** |
| --- | --- | --- | --- | --- | --- | --- | --- |
| Leuk | Low | 890 | 46.27, 7.88 | Varen (low origin) | 48 | 10 | 4.8±3.260 |
|  |  |  |  | Saxon (low origin) | 42 | 8 | 5.25±2.252 |
|  |  |  |  | Saviese (low origin) | 69 | 14 | 4.929±2.870 |
|  |  |  |  | Chäller (high origin) | 48 | 17 | 2.824±1.667 |
|  |  |  |  | Tsansfleuron (high origin) | 59 | 20 | 2.95±2.328 |
|  |  |  |  | Val de Bagne (high origin) | 82 | 25 | 3.28±2.492 |
| Zeneggen | Low | 930 | 46.31, 7.66 | Varen (low origin) | 48 | 11 | 4.364±3.641 |
|  |  |  |  | Saxon (low origin) | 38 | 7 | 5.429±0.976 |
|  |  |  |  | Saviese (low origin) | 70 | 16 | 4.375±2.446 |
|  |  |  |  | Chäller (high origin) | 54 | 16 | 3.375±1.5 |
|  |  |  |  | Tsansfleuron (high origin) | 58 | 18 | 3.222±2.045 |
|  |  |  |  | Val de Bagne (high origin) | 74 | 25 | 2.96±2.189 |
| Findeln | High | 2120 | 46.01, 7.76 | Varen (low origin) | 50 | 8 | 6.25±2.915 |
|  |  |  |  | Saxon (low origin) | 31 | 8 | 3.875±1.727 |
|  |  |  |  | Saviese (low origin) | 64 | 13 | 4.923+2.753 |
|  |  |  |  | Chäller (high origin) | 52 | 16 | 3.25±2.082 |
|  |  |  |  | Tsansfleuron (high origin) | 53 | 17 | 3.118±2.027 |
|  |  |  |  | Val de Bagne (high origin) | 88 | 26 | 3.385±2.192 |
| Oberu | High | 2150 | 46.35, 7.67 | Varen (low origin) | 43 | 10 | 4.3±2.163 |
|  |  |  |  | Saxon (low origin) | 32 | 8 | 4±1.604 |
|  |  |  |  | Saviese (low origin) | 48 | 14 | 3.429±2.277 |
|  |  |  |  | Chäller (high origin) | 66 | 19 | 3.474±2.366 |
|  |  |  |  | Tsansfleuron (high origin) | 49 | 17 | 2.882±1.996 |
|  |  |  |  | Val de Bagne (high origin) | 69 | 24 | 2.875±2.133 |

**Table S3.** Overview of the recruitment experiment of wild *D. sylvestris* populations. Transplant site name, elevational environment denominator (i.e., low or high), nr. of seeds sown per population and transplant site and nr. of maternal families and mean nr. of seeds per family (± SD) reported.

| **Transplant site** | **Transplant environment** | **Population** | **Nr. of seeds sown** | **Nr. of maternal families represented** | **Mean nr. of seeds per maternal family, ± SD** |
| --- | --- | --- | --- | --- | --- |
| Leuk | Low | Varen (low origin) | 100 | 5 | 20±0 |
|  |  | Saxon (low origin) | 100 | 5 | 20±0 |
|  |  | Saviese (low origin) | 100 | 4 | 25±4.082 |
|  |  | Chäller (high origin) | 100 | 5 | 20±0 |
|  |  | Tsansfleuron (high origin) | 100 | 4 | 25±0 |
|  |  | Val de Bagne (high origin) | 100 | 5 | 20±0 |
| Zeneggen | Low | Varen (low origin) | 100 | 5 | 20±0 |
|  |  | Saxon (low origin) | 100 | 5 | 20±0 |
|  |  | Saviese (low origin) | 100 | 5 | 20±0 |
|  |  | Chäller (high origin) | 100 | 5 | 20±0 |
|  |  | Tsansfleuron (high origin) | 100 | 3 | 33.333±23.094 |
|  |  | Val de Bagne (high origin) | 100 | 2 | 50±14.142 |
| Findeln | High | Varen (low origin) | 100 | 2 | 50±49.497 |
|  |  | Saxon (low origin) | 100 | 4 | 25±4.082 |
|  |  | Saviese (low origin) | 0 | 0 | 0 |
|  |  | Chäller (high origin) | 100 | 3 | 33.333±2.887 |
|  |  | Tsansfleuron (high origin) | 100 | 4 | 25±4.082 |
|  |  | Val de Bagne (high origin) | 100 | 4 | 25±4.082 |
| Oberu | High | Varen (low origin) | 100 | 2 | 50±49.497 |
|  |  | Saxon (low origin) | 100 | 4 | 25±4.082 |
|  |  | Saviese (low origin) | 0 | 0 | 0 |
|  |  | Chäller (high origin) | 100 | 3 | 33.333±2.887 |
|  |  | Tsansfleuron (high origin) | 100 | 4 | 25±4.082 |
|  |  | Val de Bagne (high origin) | 100 | 4 | 25±4.082 |

**Table S4.** Creation of F1 and F2 populations. Top: Sampling location of plants used to generate F1 crosses, including population name, elevational origin denominator (i.e., low or high) and nr. of grandparent plants used for F1 crosses reported. Bottom: production of F2 crosses, cage, F1 cross and nr. of plants per cross and cage reported. Note that the number after F1s refers to the different grandparent plants.

| **F0 plants used in production of F1 the generation** | | |
| --- | --- | --- |
| **Population** | **Elevational origin** | **Nr. grandparent plants** |
| Saxon | Low | 1 |
| Varen | Low | 1 |
| Tsanfleuron | High | 2 |
| **F2 crosses** | | |
| **Cage** | **F1** | **Nr. of plants** |
| 1 | Saxon_X_Tsanfleuron2 | 28 |
| 2 | Saxon_X_Tsanfleuron2 | 15 |
| 2 | Tsanfleuron_X_Varens1 | 15 |
| 3 | Tsanfleuron_X_Varens1 | 13 |
| 3 | Saxon_X_Tsanfleuron2 | 13 |
| 4 | Saxon_X_Tsanfleuron2 | 10 |
| 4 | Tsanfleuron_X_Saxon2 | 10 |
| 4 | Tsanfleuron_X_Varens1 | 10 |
| 5 | Tsansfleuron_X_Varens1 | 15 |

**Table S5.** Overview of the transplant experiment of F2 crosses. Transplant site, elevational environment denominator (i.e., low or high), cage number, number of plants per transplant size and cage and number of plants per genetic cluster as identified by PCA analysis (nrs. 1 to 3 arbitrarily chosen, Pålsson 2023, chapter 3) reported. Note that all the plant nrs. refer to plants alive at the start of 2018 and hence form the basis of our analyses.

| **Site** | **Elevational environment** | **Cage** | **Number of plants alive after transplant shock** | **Number of plants per genetic cluster, 1; 2; 3** |
| --- | --- | --- | --- | --- |
| Zeneggen | Low | 1 | 122 | 122; 0; 0 |
|  |  | 2 | 134 | 43; 67; 24 |
|  |  | 3 | 132 | 52; 56; 24 |
|  |  | 4 | 119 | 57;42;20 |
|  |  | 5 | 114 | 144; 0; 0 |
| Findeln | High | 1 | 101 | 100; 1; 0 |
|  |  | 2 | 122 | 32; 60; 30 |
|  |  | 3 | 103 | 47; 47; 9 |
|  |  | 4 | 124 | 62; 48; 14 |
|  |  | 5 | 104 | 104; 0; 0 |

| **Test** | **S_1_** | **S_2_** | | **S_3_** | **S_4_** | **S_5_** | **Standardized mean** |
| --- | --- | --- | --- | --- | --- | --- | --- |
| Model | plant height s1~ecotype*environment+(1\|site) + (1\|population/maternal family) | plant height s2~ecotype*environment+(1\|site) + (1\|population) | | plant height s3~ecotype*environment+(1\|site) + (1\|population/maternal family) | plant height s4~ecotype*environment+(1\|site) + (1\|population/maternal family) | plant height s5~ecotype*environment+(1\|site) + (1\|population/maternal family) | mean plant height~ecotype*environment+(1\|site/block) + (1\|population/maternal family) |
|  | **χ^2^; P-value** | | | | | |  |
| GxE | **7.616; 0.005** | 0.051; 0.821 | | **7.898; 0.005** | **15.921; <.0001** | **22.359; <.0001** | 0.406; 0.524 |
| *Local vs. foreign* | | | **Estimate; SE; p-value** | | | |  |
| Low env. | **-84.33; 18.5; 0.008** | **-26.2; 9.79; 0.022** | | **-88.2; 15.7; 0.003** | **-114.81;** **16.4; 0.0003** | **-** **123.1; 19.4; 0.001** | **-1.010; 0.232; 0.010** |
| High env. | **-54.20; 19.3; 0.035** | **-29.1; 11.40; 0.02** | | **-43.1; 19.4; 0.05** | **-52.04; 17.6; 0.017** | **-** **60.0; 19.2; 0.023** | **-0.935; 0.238; 0.012** |
| *Home vs. away* | | |  | | | |  |
| Low eco. | -36.41; 25.0; 0.26 | -11.7; 30.23; 0.732 | | -139.7; 61.9; 0.145 | -66.65; 35.2; 0.182 | -109.3; 33.6; 0.071 | 0.179; 0.376; 0.678 |
| High eco. | -6.28; 24.1; 0.817 | -14.6; 30.12; 0.672 | | -94.7; 61.3; 0.26 | -3.88; 34.9; 0.921 | -46.2; 33.3; 0.291 | 0.254; 0.371; 0.562 |

**Table S6.** Results of the plant height analyses. We tested for divergence in plant height by modelling it as the response variable in linear mixed effect models with a Gaussian error distribution. We tested for ecotype by environment interactions and differential performance of the ecotypes according to the local vs. foreign and home vs. away criteria of local adaptation. We tested the significance of the interaction between elevation and ecotype using likelihood ratio tests and report χ^2^ and p-values. Local vs. foreign and home vs. away contrasts were estimated using pairwise contrasts in the emmeans R package; Estimate, SE and p-values are reported, significant results are in bold. S_i_ denote the growing seasons, Low env. and Low eco. and High env. and High eco. the low and high environments and ecotypes, respectively.

| **Test** | **S_1_** | **S_2_** | | | **S_3_** | **S_4_** | **S_5_** | **Standardized mean** |
| --- | --- | --- | --- | --- | --- | --- | --- | --- |
| Model | flowering time s1~ecotype*environment+(1\|site/site_plot) + (1\|population/maternal family) | flowering time s2~ecotype*environment+(1\|site) + (1\|population/maternal family) | | | flowering time s3~ecotype*environment+(1\|site) + (1\|population/maternal family) | log(flowering time s4)~ecotype*environment+(1\|site) + (1\|population/maternal family) | flowering time s5~ecotype*environment+(1\|site) + (1\|population/maternal family) | mean flowering time~ecotype*environment+(1\|site/block) + (1\|population/maternal family) |
|  | **χ^2^; P-value** | | | | | | |  |
| GxE | 0.006; 0.940 | 0.277; 0.599 | | | 0.108; 0.743 | **11.457; <.0001** | **6.062; 0.014** | **22.842; <.0001** |
| *Local vs. foreign* | | | **Estimate; SE; p-value** | | | | |  |
| Low env. | **-26.1; 8.34; 0.034** | -12.4; 8.30; 0.202 | | | -13.8; 7.98; 0.156 | 0.901; 0.048; 0.117 | **-23.3; 9.47; 0.068** | -1.029; 0.445; 0.081 |
| High env. | **-26.0; 8.54; 0.033** | -14.9; 8.80; 0.143 | | | -15.1; 8.45; 0.131 | **0.835; 0.045; 0.023** | **-29.4; 9.51; 0.034** | **-1.480; 0.449; 0.028** |
| *Home vs. away* | | | |  | | | |  |
| Low eco. | **53.3; 12.81; 0.048** | **35.9; 7.66; 0.018** | | | **57.2; 11.66; 0.032** | **1.414; 0.102; 0.034** | **59.6; 9.94; 0.024** | 0.982; 0.571; 0.224 |
| High eco. | 53.5; 12.64; 0.051 | 33.4; 7.25; 0.031 | | | **55.9; 11.46; 0.037** | 1.309; 0.094; 0.059 | **53.4; 9.88; 0.031** | 0.531; 0.567; 0.448 |

**Table S7.** Results of the flowering time analyses. We tested for divergence in flowering time by modelling it as the response variable in linear mixed effect models with a Gaussian error distribution. We tested for environment by environment interactions and differential performance of the environments according to the local vs. foreign and home vs. away criteria of local adaptation. We tested the significance of the interaction between elevation and ecotype using likelihood ratio tests and report χ^2^ and p-values. Local vs. foreign and home vs. away contrasts were estimated using pairwise contrasts in the emmeans R package; Estimate, SE and p-values are reported, significant results are in bold. S_i_ denote the growing seasons, Low env. and Low eco. and High env. and High eco. the low and high environments and ecotypes, respectively.

**Table S8.** Results of the plant size analyses. We tested for divergence in plant size by modelling it as the response variable in linear mixed effect models with a Gaussian error distribution. We tested for ecotype by environment interactions and differential performance of the ecotypes according to the local vs. foreign and home vs. away criteria of local adaptation. We tested the significance of the interaction between elevation and ecotype using likelihood ratio tests and report χ^2^ and p-values. Local vs. foreign and home vs. away contrasts were estimated using pairwise contrasts in the emmeans R package; Estimate, SE and p-values are reported, significant results are in bold. Low env. and Low eco. and High env. and High eco. the low and high environments and ecotypes, respectively.

| **Test** | **S_1_** | **E_1_** | **S_2_** | **S_3_** | **E_3_** | **S_4_** | **E_4_** | **S_5_** | **Standardized mean** |
| --- | --- | --- | --- | --- | --- | --- | --- | --- | --- |
| Model | plant size start of s1 ~ecotype*environment+(1\|site/block)+(1\|population/maternal family) | plant size e1~ecotype*environment+(1\|site/block)+(1\|population/maternal family) | plant size s2~ecotype*environment+(1\|site/block)+(1\|population/maternal family) | plant size s3~ecotype*environment+(1\|site/block) + (1\|population/maternal family) | plant size e3~ecotype*environment+(1\|site/block) + (1\|population/maternal family) | plant size s4~ecotype*environment+(1\|site/block) + (1\|population/maternal family) | plant size e4~ecotype*environment+(1\|site/block) + (1\|population/maternal family) | plant size s5~ecotype*environment+(1\|site) + (1\|population/maternal family) | mean plant size~ecotype*environment+(1\|site/block) + (1\|population/maternal family) |
| **χ^2^; P-value** | | | | | | | | |  |
| GxE | **40.603; <.0001** | **51.014; <.0001** | **66.5; <.0001** | **49.221 ; <.0001** | **51.263; <.0001** | **40.968 ; <.0001** | **66.129; <.0001** | **41.765; <.0001** | **86.851; <.0001** |
| *Local vs. foreign* | |  |  | **Estimate; SE; P-value** | |  |  |  |  |
| Low env. | -5.52 ; 2.60; 0.093 | **-22.00; 6.19; 0.021** | **-13.00; 3.5; 0.016** | **-22.01; 6.19; 0.021** | **-26.39 ; 8.31; 0.030** | **-32.41; 7.92; 0.009** | **-43.70; 10.6; 0.010** | **-35.56; 8.46; 0.005** | **-0.736; 0.242; 0.035** |
| High env. | 2.93; 2.64; 0.319 | -3.97; 6.21; 0.554 | 1.73; 3.52; 0.646 | -3.29; 6.24; 0.623 | -1.24; 8,32; 0.888 | -5.03; 7.70; 0.545 | -2.94; 10.4; 0.789 | 5.51; 8.03; 0.522 | 0.156; 0.244; 0.555 |

| **Test** | **S_1_** | **E_1_** | **S_2_** | **S_3_** | **E_3_** | **S_4_** | **E_4_** | **S_5_** | **Standardized mean** |
| --- | --- | --- | --- | --- | --- | --- | --- | --- | --- |
| **Estimate; SE; P-value** | | | | | | | | |  |
| *Home vs. away* | |  |  |  |  |  |  |  |  |
| Low eco. | 5.45; 3.45; 0.241 | -6.37; 7.58; 0.483 | -3.8; 7.55; 0.663 | -31.95; 9.38; 0.075 | -32.56 ; 19.28; 0.230 | -35.59; 12.95; 0.103 | -56.09; 15.3; 0.061 | -21.77; 20.25; 0.389 | -0.473; 0.318; 0.268 |
| High eco. | 13.91; 3.38; 0.051 | 11.65; 7.51; 0.256 | 10.93; 7.50; 0.280 | -13.23 ; 9.30; 0.288 | -7.40; 19.23; 0.737 | -8.20; 12.96; 0.588 | -15.33; 15.3; 0.146 | 19.29; 20.21; 0.436 | 0.419; 0.315; 0.312 |

**Table S8.** Continued.

**Table S9.** Linear mixed effect models, and linear models for the effect of plant size on plant height and flowering time. We tested the significance of the interactions using likelihood ratio tests and report χ^2^ and p-values. Trends were estimated using pairwise contrasts in the emmeans R package; Response, models, estimates, SE, degrees of freedom (df), t value, and p-values are reported. S_i_ denote the growing seasons. Significant results are in bold.

| **Respone and fixed effects** | **plant size * ecotype: 2-way interaction; χ^2^; P-value** | **plant size * environment: 2-way interaction; χ^2^; P-value** | **plant size * ecotype * environment: 3-way interaction; χ^2^; P-value** | **Model** |
| --- | --- | --- | --- | --- |
| plant height s1 ~ plant size start of s1*ecotype* environment | 0.081; 0.776 | 0.271; 0.603 | 1.280;0.258 | plant height s1 ~ plant size start of s1 *ecotype*environment +(1\|site) + (1\|population/maternal family) |
| plant height s2 ~ plant size start of s2*ecotype* environment | 1.605; 0.205 | 0.000; 0.999 | -0.0062; 0.804 | plant height s2 ~ plant size start of s2*ecotype* environment+(1\|site/block) + (1\|population) |
| plant height s3 ~ plant size start of s3*ecotype* environment | 0.427; 0.514 | 5.600; 0.014 | -0.247; 0.619 | plant height s3 ~ plant size start of s3*ecotype* environment+(1\|site/block) + (1\|population/maternal family) |
| plant height s4 ~ plant size start of s4*ecotype* environment | 0.306; 0.580 | 0.623; 0.30 | 0.015; 0.902 | plant height s4 ~ plant size start of s4*ecotype* environment+(1\|site) + (1\|population/maternal family) |
| plant height s5 ~ plant size start of s5*ecotype* environment | 1.534; 0.215 | 0.381; 0.537 | 0.487; 0.485 | plant height s5 ~ plant size start of s5*ecotype*environment+(1\|site/block) + (1\|population/maternal family) |

**Table S9.** Continued.

| **Respone and fixed effects** | **plant size * ecotype: 2-way interaction; χ^2^; P-value** | **plant size * environment: 2-way interaction; χ^2^; P-value** | **plant size * ecotype * environment: 3-way interaction; χ^2^; P-value** | **Model** |
| --- | --- | --- | --- | --- |
| flowering time s1 ~ plant size start of s1*ecotype* environment | 0.373; 0.541 | **4.361; 0.037** | 0.251; 0.616 | flowering time s1 ~ plant size start of s1*ecotype*environment+(1\|site) + (1\|population/maternal family) |
| flowering time s2 ~ plant size start of s2*ecotype* environment | **4.293; 0.038** | 2.505; 0.113 | 0.191; 0.662 | flowering time s2 ~ plant size start of s2*ecotype*environment+(1\|site) + (1\|population/maternal family) |
| flowering time s3 ~ plant size start of s3*ecotype* environment | 0.711; 0.400 | 0.132; 717 | 0.004; 0.950 | flowering time s3 ~ plant size start of s3*ecotype*environment+(1\|site) + (1\|population/maternal family) |
| flowering time s4 ~ plant size start of s4*ecotype* environment | **4.86; 0.026** | 0.216; 0.642 | 0.815; 0.367 | flowering time s4 ~ plant size start of s4*ecotype*environment+(1\|site) + (1\|population/maternal family) |
| flowering time s5 ~ plant size start of s5*ecotype* environment | 3.210; 0.073 | 1.543; 0.214 | 0.234; 0.625 | flowering time s5 ~ plant size start of s5*ecotype*environment+(1\|site) + (1\|population/maternal family) |

**Table S9.** Continued.

| **Within environments** | | | |  | |  | | | |  |
| --- | --- | --- | --- | --- | --- | --- | --- | --- | --- | --- |
| **Response** | **Environment** | **Ecotype** | **Trend; plant size start of season** | | **SE** | | **df** | **t.ratio** | **p-value** | **Model** |
| plant height s1 | High environment | high | **1.324** | | **0.323** | | **104.2** | **4.097** | **0.0001** | plant height s1 ~ plant size start of s1*ecotype+(1\|site) + (1\|population) |
|  |  | low | 0.027 | | 0.842 | | 91.4 | 0.032 | 0.975 |  |
|  | Low environment | high | 1.34 | | 1.217 | | 125 | 1.101 | 0.273 | plant height s1 ~ plant size start of s1 *ecotype+(1\|site) + (1\|population/maternal family) |
|  |  | low | **1.84** | | **0.858** | | **123** | **2.145** | **0.034** |  |
| plant height s2 | High environment | high | **1.138** | | **0.389** | | **57.6** | **2.928** | **0.005** | plant height s2 ~ plant size start of s2*ecotype+(1\|site) + (1\|population/maternal family) |
|  |  | low | 0.924 | | 0.475 | | 54.8 | 1.943 | 0.057 |  |
|  | Low environment | high | 1.495 | | 0.832 | | 100 | 1.798 | 0.075 | plant height s2 ~ plant size start of s2 *ecotype) |
|  |  | low | **0.755** | | **0.292** | | **100** | **2.583** | **0.011** |  |
| plant height s3 | High environment | high | 0.692 | | 0.487 | | 63.9 | 1.421 | 0.160 | plant height s3 ~ plant size start of s3 *ecotype+(1\|site) + (1\|population/maternal family) |
|  |  | low | 1.671 | | 1.135 | | 51.4 | 1.472 | 0.147 |  |
|  | Low environment | high | **1.195** | | **0.347** | | **171** | **3.449** | **0.001** | plant height s3 ~ plant size start of s3 *ecotype+(1\|site) + (1\|population/maternal family) |
|  |  | low | **0.788** | | **0.221** | | **154** | **3.570** | **0.001** |  |
| plant height s4 | High environment | high | **1.132** | | **0.365** | | **8.28** | **3.104** | **0.014** | plant height s4 ~ plant size start of s4*ecotype+(1\|site) |
|  |  | low | 0.325 | | 0.612 | | 111.05 | 0.531 | 0.597 |  |
|  | Low environment | high | 0.753 | | 0.411 | | 98.9 | 1.831 | 0.070 | plant height s4 ~ plant size start of s4*ecotype+(1\|site) + (1\|population/maternal family) |
|  |  | low | **0.678** | | **0.174** | | **67.2** | **3.901** | **0.0002** |  |

**Table S9.** Continued.

| **Within environments** | | |  | |  | | |  |
| --- | --- | --- | --- | --- | --- | --- | --- | --- |
| **Response** | **Environment** | **Ecotype** | **Trend; plant size start of season** | **SE** | **df** | **t.ratio** | **p-value** | **Model** |
| plant height s5 | High environment | high | **0.561** | **0.103** | **188** | **5.434** | **<.0001** | plant height s5 ~ plant size start of s5 *ecotype+(1\|site) |
|  |  | low | 0.196 | 0.211 | 188 | 0.928 | 0.354 |  |
|  | Low environment | high | 0.287 | 0.458 | 102 | 0.627 | 0.32 | plant height s5 ~ plant size start of s5*ecotype+(1\|site) + (1\|population/maternal family) |
|  |  | low | 0.390 | 0.222 | 102 | 1.759 | 0.082 |  |
| flowering time s1 | High environment | high | -0.005 | 0.138 | 142 | -0.038 | 0.969 | flowering time s1 ~ plant size start of s1*ecotype + (1\|population) |
|  |  | low | -0.329 | 0.294 | 113 | -1.116 | 0.267 |  |
|  | Low environment | high | -0.292 | 0.168 | 164 | -1.741 | 0.0834 | flowering time s1 ~ plant size start of s1*ecotype + (1\|population/maternal family) |
|  |  | low | **-0.330** | **0.155** | **160** | **-2.126** | **0.035** |  |
| flowering time s2 | High environment | high | -0.195 | 0.097 | 35.8 | -2.012 | 0.052 | flowering time s2 ~ plant size start of s2*ecotype + (1\|population) |
|  |  | low | **-0.917** | **0.231** | **35.6** | **-3.969** | **0.0003** |  |
|  | Low environment | high | -0.004 | 1.637 | 31 | -0.002 | 0.998 | flowering time s2 ~ plant size start of s2*ecotype + (1\|population) |
|  |  | low | -0.284 | 0.247 | 32.7 | -1.152 | 0.258 |  |
| flowering time s3 | High environment | high | **-0.522** | **0.112** | **71.2** | **-4.650** | **<.0001** | flowering time s3 ~ plant size start of s3*ecotype + (1\|population) |
|  |  | low | -0.457 | 0.412 | 61.1 | -1.111 | 0.271 |  |
|  | Low environment | high | -0.072 | 0.99 | 172 | -0.726 | 0.469 | flowering time s3 ~ plant size start of s3*ecotype + (1\|population) |
|  |  | low | **-0.165** | **0.063** | **174** | **-2.631** | **0.009** |  |

| **Within environments** | | |  | |  | | |  |
| --- | --- | --- | --- | --- | --- | --- | --- | --- |
| **Response** | **Environment** | **Ecotype** | **Trend; plant size start of season** | **SE** | **df** | **t.ratio** | **p-value** | **Model** |
| flowering time s4 | High environment | high | -0.209 | 0.095 | 107 | -2.207 | 0.029 | flowering time s4 ~ plant size start of s4*ecotype +(1\|site/block) + (1\|population) |
|  |  | low | 0.222 | 0.202 | 112 | 1.100 | 0.274 |  |
|  | Low environment | high | -0.189 | 0.111 | 110 | -1.697 | 0.092 | flowering time s4 ~ plant size start of s4*ecotype + (1\|population) |
|  |  | low | -0.052 | 0.047 | 110 | -1.095 | 0.276 |  |
| flowering time s5 | High environment | high | 0.001 | 0.028 | 216 | 3.539 | 0.001 | flowering time s5 ~ plant size start of s5*ecotype + |
|  |  | low | -0.077 | 0.056 | 216 | -1.393 | 0.165 |  |
|  | Low environment | high | -0.209 | 0.076 | 165 | -2.738 | 0.007 | flowering time s5 ~ plant size start of s5*ecotype+(1\|population) |
|  |  | low | **-0.220** | **0.038** | **165** | **-5.751** | **<.0001** |  |
| **Response** | **Environment** | **Ecotype** | **ExT (trait) interaction, χ^2^; P-value** | **Trend; plant size start of season** | **SE** | **df** | **z value** | **p-value** |
| mean flowering time | High environment | high | **8.629; 0.003** | -0.069 | 0.059 | 279 | -1.161 | 0.247 |
|  |  | low |  | **-0.459** | **0.120** | **279** | **-3.829** | **0.0002** |
|  | Low environment | high | 1.058, 0.304 | **-0.217** | **0.088** | **341** | **-2.473** | **0.014** |
|  |  | low |  | **-0.330** | **0.067** | **342** | **-4.928** | **<.0001** |
| mean height | High environment | high | 2.744; 0.098 | **0.494** | **0.070** | **291** | **7.084** | **<.0001** |
|  |  | low |  | **0.277** | **0.116** | **291** | **2.390** | **0.0097** |
|  | Low environment | high | **3.930; 0.003** | **0.586** | **0.105** | **312** | **5.600** | **<.0001** |
|  |  | low |  | **0.210** | **0.070** | **312** | **2.990** | **0.003** |

**Table S9.** Continued.

**Table S9**. Continued.

| **Respone and fixed effects** | **plant size * ecotype: 2-way interaction; χ^2^; P-value** | **plant size *environment: 2-way interaction; χ^2^; P-value** | **plant size * ecotype * environment: 3-way interaction; χ^2^; P-value** | **Model** |
| --- | --- | --- | --- | --- |
| mean flowering time ~ mean size * ecotype *environment | 0.626; 0.429 | **4.281; 0.039** | 0.112; 0.738 | mean flowering time ~ mean plant size*ecotype*environment+(1\|site/block) + (1\|population/maternal family) |
| mean height ~ mean size * ecotype *environment | 1.491; 0.222 | 0.029; 0.864. | 0.012; 0.911 | mean height ~ mean plant size*ecotype*environment+(1\|site/block) + (1\|population/maternal family) |

**Table S10**. Results of the LTRE showing the contribution of the vital rates of the foreign ecotype to population growth rate in the two transplant environments. Survival and reproductive vital rates throughout the life cycle are indicated by T and R, respectively. Mean estimates based on 20 000 bootstrap replicated are reported.

| **Environment** | **T_1_** | **R_1_** | **T_2_** | **T_3_** | **R_2_** | **T_4_** | **T_5_** | **R_3_** | **T_6_** | **T_7_** | **R_4_** | **T_8_** | **T_9_** | **R_5_** | **T_10_** |
| --- | --- | --- | --- | --- | --- | --- | --- | --- | --- | --- | --- | --- | --- | --- | --- |
| High | -0.052 | -0.006 | -0.003 | -0.008 | 0.002 | --0.012 | -0.007 | -0.008 | -0.002 | -0.015 | -0.059 | -2.43E-04 | 1.89E-04 | -0.071 | 0 |
| Low | -0.005 | 0.007 | -0.027 | -0.012 | -0.005 | -0.009 | -0.004 | -0.212 | -0.006 | -0.015 | -0.005 | -0.001 | 0.001 | -0.081 | 0 |

**Table S11**. Results of the cumulative survival analyses. We modelled survival throughout the experiment using cox proportional hazard models and tested for ecotype by environment interactions and differential performance of the ecotypes according to the local vs. foreign and home vs. away criteria of local adaptation. We report coefficients, hazard ratios and p-values, significant results are in bold. Low env. and High env. denote low and high environment, respectively.

| **Test** | **Coefficients** | **Hazard ratio** | **p-values** |
| --- | --- | --- | --- |
| GxE | **-2.457** | **0.086** | **<.0001** |
| *Local vs. foreign* | |  |  |
| Low env. | **-0.734** | **0.48** | **0.002** |
| High env. | **1.627** | **5.088** | **<.0001** |
| *Home vs. away* | |  |  |
| Low ecotype | -0.587 | 0.556 | 0.066 |
| High ecotype | **1.96** | **7.099** | **0.016** |

**Table S12**. Results of survival probability analyses. We tested for adaptation in survival probability by modelling it as the response variable in generalized linear mixed effect models with binomial error distribution. We tested for eccotype by environment interactions and differential performance of the ecotypes according to the local vs. foreign and home vs. away criteria of local adaptation. We tested the significance of the interaction between elevation and ecotype using likelihood ratio tests and report χ^2^ and p-values. Local vs. foreign and home vs. away contrasts were estimated using pairwise contrasts in the emmeans R package; Models, estimate, SE and p-values are reported, significant results are in bold. W_i_ and S_i_ denote the season. Low env. and Low eco. and High env. and High eco. the low and high environments and ecotypes, respectively.

| **Test** | **W_1_** | **S_1_** | **W_2_** | **S_2_** | **W_3_** | **S_3_** | **W_4_** | **S_4_** | **W_5_** |
| --- | --- | --- | --- | --- | --- | --- | --- | --- | --- |
| Model | survival probability w1 (y/n) ~ecotype*environment+(1\|site/block) | survival probability s1 (y/n)~ecotype*environment+(1\|site/block) + (1\|population/maternal family) | survival probability w2 (y/n)~ecotype*environment+(1\|site/block) | survival probability s2 (y/n)~ecotype*environment+(1\|site/block) | NA | survival probability s3 (y/n)~ecotype*environment+(1\|site/block) | survival probability w4 (y/n)~ecotype*environment+(1\|site/block)+ (1\|population/maternal family) | NA | NA |
| **χ^2^; P-value** | | | | | | | | | |
| GxE | **62.668; <.0001** | **7.610; 0.006** | **17.01; <.0001** | **16.795; <.0001** | NA | **4.674; 0.031** | **42.559; <.0001** | NA | NA |
|  |  |  |  |  |  |  |  | NA | NA |
| *Local vs. foreign* | |  | **Estimate; SE; P-value** | |  |  |  | NA | NA |
| Low env. | 0.76; 0.191; 0.2743 | **0.436; 0.132; 0.0062** | **3.878; 1.650; 0.0015** | 0.444; 0.188; 0.0547 | NA | **0.380; 0.121; 0.003** | 0.804; 1.089; 0.872 | NA | NA |
| High env. | **11.814; 3.017; <.0001** | 1.805; 0.839; 0.2034 | **0.339; 0.148; 0.0134** | **4.613; 1.779; 0.0001** | NA | 1.72; 0.997; 0.353 | **6.492; 2.832; <.0001** | NA | NA |
| *Home vs. away* | |  |  |  |  |  |  | NA | NA |
| Low eco. | **0.103; 0.103; 0.023** | 3.928; 2.897; 0.0636 | **0.242; 0.146; 0.019** | **0.236; 0.133; 0.010** | NA | 2.01; 1.329; 0.291 | 0.804; 1.089; 0.872 | NA | NA |
| High eco. | 1.604; 1.612; 0.6384 | **16.275; 11.609; 0.0001** | 2.760; 1.513; 0.0640 | 2.450; 1.304; 0.092 | NA | **9.07; 5.068; 0.0001** | **24.266; 33.034; 0.019** | NA | NA |

| **Test** | **S_1_** | **S_2_** | **S_3_** | **S_4_** | **S_5_** | **Cumulative seed count** |
| --- | --- | --- | --- | --- | --- | --- |
| Model | seed count s1~ecotype*environment+(1\|site/block) + (1\|population/maternal family) | seed count s2~ecotype*environment+(1\|site/block) + (1\|population) | seed count s3~ecotype*environment+(1\|site/block) + (1\|population) | seed count s4~ecotype*environment+(1\|site/block) + (1\|population) | seed count s5~ecotype*environment+(1\|site/block) + (1\|population) | cumulative seed count~ecotype+environment+(1\|site/block) + (1\|population/maternal family) |
| **χ^2^; P-value** | | | | | | |
| GxE | **10.349; 0.001** | **103.218; <.0001** | **319.57; <.0001** | **92.661; <.0001** | **704.72; <.0001** | **3262.9; <.0001** |
| *Local vs. foreign* | |  | **Estimate; SE; P-value** | |  |  |
| Low env. | 1.105; 0.356; 0.757 | 1.85; 1.0; 0.256 | **0.311; 0.101; <0.001** | **0.31; 0.084; <.0001** | **0.279; 0.138; 0.010** | **0.214; 0.061; <.0001** |
| High env. | 0.350; 0.159; 0.021 | **0.33; 0.17; 0.028** | 0.832; 0.274; 0.577 | **1.94; 0.295; <.0001** | 0.892; 0.441; 0.817 | 0.982; 0.281; 0.951 |
| *Home vs. away* | |  |  |  |  |  |
| Low eco. | 0.391; 0.564; 0.515 | **11.27; 3.56; <.0001** | 0.094; 0.103; 0.031 | 0.90; 0.227; 0.663 | **0.169; 0.050; <.0001** | **0.250; 0.067; <.0001** |
| High eco. | 0.124; 0.175; 0.139 | 2.03; 0.63; 0.023 | 0.252; 0.275; 0.207 | **5.69; 1.927; <.0001** | **0.541; 0.160; 0.038** | 1.146; 0.305; 0.608 |

**Table S13**. Results of seed count analyses. We tested for adaptation in seed count by modelling it as the response variable in generalized linear mixed effect models with zero inflated poisson error distribution. We tested for ecotype by environment interactions and differential performance of the ecotypes according to the local vs. foreign and home vs. away criteria of local adaptation. We tested the significance of the interaction between elevation and ecotype using likelihood ratio tests and report χ^2^ and p-values. Local vs. foreign and home vs. away contrasts were estimated using pairwise contrasts in the emmeans R package; Models, estimate, SE and p-values are reported, significant results are in bold. S_i_ denote the growing season. Low env. and Low eco. and High env. and High eco. the low and high environments and ecotypes, respectively.

**Table S14.** Results of flowering probability analyses. We tested for adaptation in flowering probability by modelling it as the response in generalized linear mixed effect models with binomial error distribution. We tested for ecotype by environment interactions and differential performance of the ecotypes according to the local vs. foreign and home vs. away criteria of local adaptation. We tested the significance of the interaction between elevation and ecotype using likelihood ratio tests and report χ^2^ and p-values. Local vs. foreign and home vs. away contrasts were estimated using pairwise contrasts in the emmeans R package; Models, estimate, SE and p-values are reported, significant results are in bold. S_i_ denote the growing season. Low env. and Low eco. and High env. and High eco. the low and high environments and ecotypes, respectively.

| **Test** | **S_1_** | **S_2_** | **S_3_** | **S_4_** | **S_5_** |
| --- | --- | --- | --- | --- | --- |
| Model | flowering probability s1 (y/n) ~ecotype*environment +(1\|site) | flowering probability s2 (y/n) ~ecotype* environment+(1\|site/block) + (1\|population/maternal family) | flowering probability s3 (y/n) ~ecotype*environment+(1\|site/block) + (1\|population/maternal family) | flowering probability s4 (y/n)~ecotype*environment+(1\|site/block) + (1\|population/maternal family) | flowering probability s5 (y/n)~ecotype*environment+(1\|site/block) + (1\|population/maternal family) |
| **χ^2^; P-value** | | | | | |
| GxE | **6.298; 0.012** | **13.264; <.0001** | **5.444; 0.012** | **17.242; <.0001** | **11.145; <.0001** |
| *Local vs. foreign* | |  | **Estimate; SE; P-value** |  |  |
| Low env. | **1.716; 0.308; 0.003** | **0.253;0.1063; 0.0011** | **0.425; 0.163; 0.026** | **0.202; 0.105; 0.002** | 0.643; 0.297; 0.339 |
| High env. | **3.722;0.968; <.0001** | 1.027; 0.460; 0.953 | 1.009; 0.433; 0.983 | 1.734; 0.726; 0.188 | **2.839; 1.208; 0.014** |
| *Home vs. away* | |  |  |  |  |
| Low eco. | **0.402; 0.113; 0.001** | **0.227; 0.097; 0.001** | **0.116; 0.054; <.0001** | 0.119; 0.136; 0.062 | 0.247; 0.183; 0.059 |
| High eco. | 0.872; 0.152; 0.432 | 0.922; 0.369 ; 0.839 | **0.275; 0.115; 0.002** | 1.734; 0.726; 0.188 | 1.093; 0.797; 0.9032 |

| **Env.jjjjjj** | **Eco.nnn** | **T_1_** | **R_1_** | **T_2_** | **T_3_** | **R_2_** | **T_4_** | **T_5_** | **R_3_** | **T_6_** | **T_7_** | **R_4_** | **T_8_** | **T_9_** | **R_5_** | **T_10_** |
| --- | --- | --- | --- | --- | --- | --- | --- | --- | --- | --- | --- | --- | --- | --- | --- | --- |
| High | High | 0.121 (0.115, 0.126) | 0.007 (0.004, 0.012) | 0.113 (0.109, 0.118) | 0.113 (0.109, 0.118) | 0.003 (0.001, 0.006) | 0.111 (0.107, 0.115) | 0.111 (0.107, 0.115) | 0.01 (0.005, 0.019) | 0.1 (0.094, 0.106) | 0.1 (0.094, 0.106) | 0.046 (0.03, 0.063) | 0.055 (0.041, 0.068) | 0.055 (0.041, 0.068) | 0.055 (0.041, 0.068) | 0 (0.041, 0.068) |
| High | Low | 0.111 (0.105, 0.123) | 0 (0, 0.001) | 0.111 (0.105, 0.123) | 0.111 (0.105, 0.123) | 0.007 (0.002, 0.034) | 0.103 (0.095, 0.109) | 0.103 (0.095, 0.109) | 0.009 (0.002, 0.025) | 0.095 (0.084, 0.101) | 0.095 (0.084, 0.101) | 0.015 (0.004, 0.037) | 0.08 (0.062, 0.092) | 0.08 (0.062, 0.092) | 0.08 (0.062, 0.092) | 0 (0.062, 0.092) |
| Low | High | 0.166 (0.141, 0.193) | 0.036 (0.019, 0.074) | 0.13 (0.113, 0.146) | 0.13 (0.113, 0.146) | 0.001 (0, 0.005) | 0.129 (0.112, 0.145) | 0.129 (0.112, 0.145) | 0.092 (0.048, 0.128) | 0.037 (0.016, 0.065) | 0.037 (0.016, 0.065) | 0.001 (0, 0.003) | 0.037 (0.015, 0.065) | 0.037 (0.015, 0.065) | 0.037 (0.015, 0.065) | 0 (0.015, 0.065) |
| Low | Low | 0.154 (0.144, 0.164) | 0.012 (0.006, 0.022) | 0.143 (0.133, 0.151) | 0.143 (0.133, 0.151) | 0.003 (0.001, 0.006) | 0.14 (0.13, 0.148) | 0.14 (0.13, 0.148) | 0.107 (0.083, 0.127) | 0.033 (0.021, 0.047) | 0.033 (0.021, 0.047) | 0.002 (0.001, 0.004) | 0.031 (0.019, 0.045) | 0.031 (0.019, 0.045) | 0.031 (0.019, 0.045) | 0 (0.019, 0.045) |

**Table S15.** Influence of specific vital rates on population growth rate at the two transplant environments. Influence of specific vital rates on population growth rates extracted from the matrix population models of the elevational ecotypes growing in the low and high environments expressed as elasticities. Survival and reproductive vital rates throughout the life cycle are indicated by T and R, respectively, Mean values are based on 20 000 bootstrap replicates and 95% bias corrected confidence intervals are reported. Env. and Eco. denote the environments and the ecotypes, respectively.

**Table S16.** Stable age distribution. Stable age distribution of the elevational ecotypes growing in the low and high environments. Winter and summer stages are represented by W and S, respectively. Values are based on 20 000 bootstrap replicates and 95% bias corrected confidence intervals are reported. Env. and Eco. denote the environments and the ecotypes, respectively.

| **Env.** | **Eco.** | **W_1_** | **S_1_** | **W_2_** | **S_2_** | **W_3_** | **S_3_** | **W_4_** | **S_4_** | **W_5_** | **S_5_** |
| --- | --- | --- | --- | --- | --- | --- | --- | --- | --- | --- | --- |
| High | High | 0.15 (0.132, 0.17) | 0.129 (0.113, 0.149) | 0.116 (0.102, 0.131) | 0.104 (0.094, 0.116) | 0.093 (0.084, 0.103) | 0.086 (0.077, 0.096) | 0.078 (0.068, 0.088) | 0.069 (0.06, 0.079) | 0.063 (0.052, 0.074) | 0.058 (0.049, 0.069) |
| High | Low | 0.074 (0.05, 0.105) | 0.053 (0.036, 0.074) | 0.06 (0.045, 0.081) | 0.066 (0.051, 0.085) | 0.069 (0.056, 0.087) | 0.08 (0.066, 0.098) | 0.093 (0.077, 0.113) | 0.094 (0.075, 0.116) | 0.111 (0.086, 0.14) | 0.136 (0.107, 0.173) |
| Low | High | 0.217 (0.176, 0.266) | 0.176 (0.147, 0.211) | 0.122 (0.106, 0.141) | 0.106 (0.094, 0.122) | 0.094 (0.08, 0.109) | 0.054 (0.045, 0.064) | 0.068 (0.051, 0.087) | 0.036 (0.024, 0.049) | 0.032 (0.02, 0.047) | 0.031 (0.018, 0.046) |
| Low | Low | 0.413 (0.381, 0.447) | 0.237 (0.212, 0.264) | 0.131 (0.115, 0.148) | 0.084 (0.072, 0.097) | 0.054 (0.045, 0.064) | 0.054 (0.045, 0.064) | 0.022 (0.016, 0.027) | 0.011 (0.008, 0.014) | 0.007 (0.005, 0.009) | 0.004 (0.003, 0.006) |

**Table S17.** Generalized linear mixed effect models, and generalized linear models for the effect of plant size on flowering probability. We tested the significance of the interactions using likelihood ratio tests and report χ^2^ and p-values. Trends were estimated using pairwise contrasts in the emmeans R package; Response, models, estimates, SE, degrees of freedom (df), z value, and p-values are reported. S_i_ denote the growing seasons. Significant results are in bold.

| **Respone and fixed effects** | **plant size*ecotype: 2-way interaction; χ^2^; P-value** | **plant size*environment: 2-way interaction; χ^2^; P-value** | **plant size * ecotype * environment: 3-way interaction; χ^2^; P-value** | **Model** |
| --- | --- | --- | --- | --- |
| flowering probability s1 ~ plant size start of s1 *ecotype* environment | **13.12; 0.0003** | 0.00; 0.99 | 0.15; 0.697 | flowering probability s1 (y/n) ~ plant size start of s1 ecotype*environment+(1\|site) + (1\|population/maternal family) |
| flowering probability s2 ~ plant size start of s2*ecotype* environment | 1.13; 0.288 | 0.01; 0.936 | 0.63; 0.427 | flowering probability s2 (y/n) ~ plant size start of s2*ecotype*environment) |
| flowering probability s3 ~ plant size start of s3*ecotype* environment | 0.0; 0.937 | 2.1; 0.149 | 0.4; 0.536 | flowering probability s3 (y/n) ~ plant size start of s3*ecotype*environment |
| flowering probability s4 ~ plant size start of s4*ecotype* environment | 0.1; 0.736 | **4.4; 0.036** | 0.8; 0.375 | flowering probability s4 (y/n) ~ plant size start of s4*ecotype*environment |

**Table S17.** Continued.

| **Respone and fixed effects** | | **plant size*ecotype: 2-way interaction; χ^2^; P-value** | | **plant size*environment: 2-way interaction; χ^2^; P-value** | **plant size * ecotype * environment: 3-way interaction; χ^2^; P-value** | | | | **Model** | | | |
| --- | --- | --- | --- | --- | --- | --- | --- | --- | --- | --- | --- | --- |
| flowering probability s5 ~ plant size start of s5*ecotype* environment | | 0.99; 0.319 | | -8.34; 0.004 | 2.52; 0.112 | | | | flowering probability s5 (y/n) ~ plant size start of s5*ecotype*environment+(1\|site/block) + (1\|population) | | | |
| flowering probability ~ plant size *ecotype* environment + year | | 3.12; 0.077 | | 0.01; 0.942 | **6.54;** **0.011** | | | | flowering probability ~ plant size*ecotype*environment+year+(1\|site/block)+(1\|population) | | | |
| **Within environments** | | |  |  | |  |  |  | |  |  |  |
| **Response** | **Environment** | | **Ecotype** | **ExT (trait) interaction, χ^2^; P-value** | | **Trend; plant size start of season** | **SE** | **df** | | **z value** | **p-value** | **Model** |
| flowering probability s1 | High environment | | high | **4.4; 0.036** | | **0.136** | **0.017** | **Inf** | | **8.200** | **<.0001** | flowering probability s1 (y/n) ~ plant size start of s1*ecotype + (1\|population) |
|  |  |  | low |  |  | **0.076** | **0.023** | **Inf** | | **3.340** | **0.0008** |  |
|  | Low environment | | high | **11.6; 0.001** | | **0.176** | **0.027** | **Inf** | | **6.510** | **<.0001** | flowering probability s1 (y/n) ~ plant size start of s1*ecotype+(1\|site/block) + (1\|population/maternal family) |
|  |  |  | low |  |  | **0.078** | **0.018** | **Inf** | | **4.390** | **<.0001** |  |
| flowering probability s2 | High environment | | high | 0.03; 0.85 | | **0.065** | **0.012** | **Inf** | | **5.390** | **<.0001** | flowering probability s2 (y/n) ~ plant size start of s2*ecotype + (1\|population/maternal family) |
|  |  |  | low |  |  | **0.069** | **0.018** | **Inf** | | **3.810** | **0.0001** |  |
|  | Low environment | | high | 1.75; 0.19 | | **0.124** | **0.028** | **Inf** | | **4.5** | **<.0001** | flowering probability s2 (y/n) ~ plant size start of s2*ecotype+(1\|site/block) + (1\|population/maternal family) |
|  |  |  | low |  |  | **0.085** | **0.016** | **Inf** | | **5.330** | **<.0001** |  |

| **Within environments** | |  |  |  |  |  |  |  |  |
| --- | --- | --- | --- | --- | --- | --- | --- | --- | --- |
| **Response** | **Environment** | **Ecotype** | **ExT (trait) interaction, χ^2^; P-value** | **Trend; plant size start of season** | **SE** | **df** | **z value** | **p-value** | **Model** |
| flowering probability s3 | High environment | high | 0.53; 0.47 | **0.063** | **0.011** | **Inf** | **5.690** | **<.0001** | flowering probability s3 (y/n) ~ plant size start of s3*ecotype + (1\|population) |
|  |  | low |  | **0.081** | **0.024** | **Inf** | **3.400** | **0.0007** |  |
|  | Low environment | high | 0.22; 0.64 | **0.0460** | **0.011** | **Inf** | **4.310** | **<.0001** | flowering probability s3 (y/n) ~ plant size start of s3*ecotype + (1\|population) |
|  |  | low |  | **0.040** | **0.009** | **Inf** | **4.650** | **<.0001** |  |
| Flowering probability s4 | High environment | high | 0.8; 0.373 | **0.064** | **0.009** | **Inf** | **6.930** | **<.0001** | flowering probability s4 (y/n) ~ plant size start of s4*ecotype |
|  |  | low |  | **0.083** | **0.020** | **Inf** | **4.110** | **<.0001** |  |
|  | Low environment | high | 0.1; 0.74 | **0.046** | **0.016** | **Inf** | **2.810** | **0.0050** | flowering probability s4 (y/n) ~ plant size start of s4*ecotype |
|  |  | low |  | **0.040** | **0.009** | **Inf** | **4.440** | **<.0001** |  |
| Flowering probability s5 | High environment | high | 3.09; 0.079 | **0.013** | **0.004** | **Inf** | **3.000** | **0.003** | flowering probability s5 (y/n) ~ plant size start of s5*ecotype+(1\|site) + (1\|population) |
|  |  | low |  | **0.029** | **0.008** | **Inf** | **3.460** | **0.0005** |  |
|  | Low environment | high | 0.88; 0.35 | 0.002 | 0.011 | Inf | 0.2 | 0.841 | flowering probability s5 (y/n) ~ plant size start of s5*ecotype+(1\|site) + (1\|population) |
|  |  | low |  | -0.009 | 0.007 | Inf | -1.22 | 0.223 |  |
| flowering probability | High environment | high | 0.95; 0.33 | **0.908** | **0.073** | **Inf** | **12.370** | **<.0001** | flowering probability ~ plant size*ecotype+year+(1\|site/block) + (1\|population/maternal family) |
|  |  | low |  | **1.048** | **0.126** | **Inf** | **8.320** | **<.0001** |  |
|  | Low environment | high | **15.3; <.0001** | **1.256** | **0.135** | **Inf** | **9.300** | **<.0001** | flowering probability ~ plant size*ecotype+year+(1\|site/block) + (1\|population/maternal family) |
|  |  | low |  | **0.663** | **0.082** | **Inf** | **8.090** | **<.0001** |  |

**Table S17.** Continued.

**Table S18**. Generalized linear mixed effect models, and generalized linear models for the effect of plant size on survival probability. S_i_ and W_i_ indicate summer and winter stages, respectively, s_i_ and e_i_ denote start and end of growing seasons, respectively. We tested the significance of the interactions using likelihood ratio tests and report χ^2^ and p-values. Trends were estimated using pairwise contrasts in the emmeans R package; Response, models, estimates, SE, degrees of freedom (df), z value, and p-values are reported. Significant results are in bold.

| **Respone and fixed effects** | **plant size*ecotype: 2-way interaction; χ^2^; P-value** | **plant size*environment: 2-way interaction; χ^2^; P-value** | **plant size size * ecotype * environment: 3-way interaction; χ^2^; P-value** | **Model** |
| --- | --- | --- | --- | --- |
| survival probability S1 ~ plant size s1 *ecotype* environment | 2.50; 0.114 | 1.15; 0.285 | 0.21; 0.643 | survival S1 (y/n) ~ plant size s1*ecotype*environment+ (1\|site) |
| survival probability W2 ~ plant size e1 *ecotype* environment | 0.17; 0.677 | 2.43; 0.119 | **6.77; 0.009** | survival W2 (y/n) ~ plant size e1*ecotype*environment+ (1\|site) |
| survival probability S2 ~ plant size s2 *ecotype* environment | 2.13; 0.145 | 2.72; 0.099 | 3.34; 0.068 | survival S2 (y/n) ~ plant size s2*ecotype*environment |
| survival probability W3 ~ plant size s2 *ecotype* environment | NA | NA | NA | NA |
| survival probability S3 ~ plant size s3 *ecotype* environment | 2.0; 0.161 | 3.6; 0.057 | 0.6; 0.457 | survival S3 (y/n) ~ plant size s3*ecotype*environment |
| survival probability W4 ~ plant size e3 *ecotype* environment | 0.9; 0.351 | 3.5; 0.062 | **8.3; 0.004** | survival W4 (y/n) ~ plant size e3*ecotype*environment |
| survival probability S4 ~ plant size s4 *ecotype* environment | NA | NA | NA | NA |
| survival probability W5 ~ plant size e4 *ecotype* environment | NA | NA | NA | NA |
| survival probability S5 ~ plant size s5 *ecotype* environment | NA | NA | NA | NA |
| survival probability ~ plant size * ecotype * environment | **4.83; 0.028** | **3.95; 0.047** | **8.54; 0.004** | survival probability ~ plant size*ecotype*environment+year+(1\|site)+(1\|population) |

**Table S18.** Continued.

| **Within environments** | |  |  |  |  |  |  |  |  |
| --- | --- | --- | --- | --- | --- | --- | --- | --- | --- |
| **Response** | **Environment** | **Ecotype** | **ExT (trait) interaction, χ^2^; P-value** | **Trend; plant size start of season** | **SE** | **df** | **z value** | **p-value** | **Model** |
| survival probability S1 | High environment | high | 0.36; 0.55 | **0.067** | **0.027** | **Inf** | **2.436** | **0.015** | survival S1 (y/n) ~ plant size s1*ecotype + (1\|site) |
|  |  | low |  | 0.041 | 0.032 | Inf | 1.271 | 0.204 |  |
|  | Low environment | high | 2.42; 0.12 | **0.094** | **0.0237** | **Inf** | **3.950** | **0.0001** | survival S1 (y/n) ~ plant size s1*ecotype+(1\|site/block) |
|  |  | low |  | **0.045** | **0.022** | **Inf** | **2.070** | **0.0389** |  |
| survival probability W2 | High environment | high | 3.67; 0.055 | **0.0837** | **0.0273** | **Inf** | **3.062** | **0.0022** | survival W2 (y/n) ~ plant size e1*ecotype + (1\|site/block) |
|  |  | low |  | 0.0268 | 0.0157 | Inf | 1.711 | 0.0872 |  |
|  | Low environment | high | 3.76; 0.052 | -0.014 | 0.014 | Inf | -1.015 | 0.310 | survival W2 (y/n) ~ plant size e1*ecotype+(1\|site/block) |
|  |  | low |  | 0.029 | 0.018 | Inf | 1.587 | 0.113 |  |
| survival probability S2 | High environment | high | 5.38; 0.02 | 0.157 | 0.046 | Inf | 1.344 | 0.0006 | survival S2 (y/n) ~ plant size s2*ecotype |
|  |  | low |  | **0.050** | **0.022** | **Inf** | **2.270** | **0.0233** |  |
|  | Low environment | high | 0.084; 0.77 | 0.025 | 0.033 | Inf | 0.778 | 0.437 | survival S2 (y/n)~ plant size s2*ecotype |
|  |  | low |  | 0.038 | 0.030 | Inf | 1.266 | 0.206 |  |
| survival probability W3 | High environment | high | NA | NA | NA | NA | NA | NA | NA |
|  |  | low |  | NA | NA | NA | NA | NA |  |
|  | Low environment | high | NA | NA | NA | NA | NA | NA | NA |
|  |  | low |  | NA | NA | NA | NA | NA |  |
| survival probability S3 | High environment | high | 0.16; 0.692 | 0.079 | 0.035 | Inf | 2.251 | 0.024 | survival S3 (y/n) ~ plant size s3*ecotype |
|  |  | low |  | 0.108 | 0.066 | Inf | 1.635 | 0.102 |  |
|  | Low environment | high | 2.25; 0.13 | **0.036** | **0.0131** | **Inf** | **2.739** | **0.006** | survival S3 (y/n) ~ plant size s3*ecotype+(1\|site) |
|  |  | low |  | 0.010 | 0.0110 | Inf | 0.936 | 0.349 |  |

| **Within environments** | |  |  |  |  |  |  |  |  |
| --- | --- | --- | --- | --- | --- | --- | --- | --- | --- |
| **Response** | **Environment** | **Ecotype** | **ExT (trait) interaction, χ^2^; P-value** | **Trend; plant size start of season** | **SE** | **df** | **z value** | **p-value** | **Model** |
| survival probability W4 | High environment | high | **9.04; 0.003** | **0.012** | **0.031** | **Inf** | **3.890** | **0.0001** | survival W4 (y/n) ~ plant size e3*ecotype |
|  |  | low |  | **0.034** | **0.014** | **Inf** | **2.430** | **0.015** |  |
|  | Low environment | high | 0.01; 0.91 | 0.020 | 0.012 | Inf | 1.715 | 0.086 | survival W4 (y/n) ~ plant size e3*ecotype+(1\|site/block)+(1\|population) |
|  |  | low |  | **0.022** | **0.010** | **Inf** | **2.223** | **0.026** |  |
| survival probability S4 | High environment | high | NA | NA | NA | NA | NA | NA |  |
|  |  | low |  | NA | NA | NA | NA | NA |  |
|  | Low environment | high | NA | NA | NA | NA | NA | NA |  |
|  |  | low |  | NA | NA | NA | NA | NA |  |
| survival probability | High environment | high | **13.7; 0.0002** | **1.525** | **0.190** | **Inf** | **8.030** | **<.0001** | survival probability ~ plant size*ecotype+year+(1\|site) + (1\|population) |
|  |  | low |  | **0.739** | **0.129** | **Inf** | **5.750** | **<.0001** |  |
|  | Low environment | high | 0.02; 0.89 | **0.783** | **0.119** | **Inf** | **6.569** | **<.0001** | survival probability ~ plant size*ecotype+year+(1\|site/block) + (1\|population) |
|  |  | low |  | **0.806** | **0.109** | **Inf** | **7.372** | **<.0001** |  |

**Table S18.** Continued.

| **Respone and fixed effects** | **plant size size*ecotype: 2-way interaction; χ^2^; P-value** | **plant size size*environment: 2-way interaction; χ^2^; P-value** | **plant size size*ecotype * environment: 3-way interaction; χ^2^; P-value** | **Model** |
| --- | --- | --- | --- | --- |
| seed count S1 ~ plant size s1 *ecotype* environment | **110.04; <.0001** | **42.20; <.0001** | **11.51; <.0001** | seed count S1 ~ plant size s1*altitude*environment+(1\|site/block) |
| seed count S2 ~ plant size s2 *ecotype* environment | **41.53; <.0001** | **58.78; <0.0001** | **5.42; 0.02** | seed count S2 ~ plant size s2*altitude* environment +(1\|site/block) + (1\|population) |
| seed count S3 ~ plant size s3 *ecotype* environment | **16.49; <.0001** | 2.61; 0.106 | 0.12; 0.725 | seed count S3 ~ plant size s3*altitude*  environment+(1\|site/block)+(1\|population/maternal family) |
| seed count S4 ~ plant size s4 *ecotype* environment | NA | NA | NA | NA |
| seed count S5 ~ plant size s5 *ecotype* environment | **25.90; <.0001** | **3.94; 0.047** | **25.06; <.0001** | seed count S5 ~ plant size s5*altitude* environment+(1\|site/block)+(1\|population) |
| seed count ~ plant size size * ecotype * Environment | 0.00; 0.94 | 0.01; 0.90 | 0.51; 0.47 | mean seed ~ mean size*altitude* environment+(1\|site/block) + (1\|population) |

**Table S19.** Generalized linear mixed effect models, and generalized linear models for the effect of plant size on seed count. S_i_ indicate stage, and s_i_ the start of the growing season. We tested the significance of the interactions using likelihood ratio tests and report χ^2^ and p-values. Trends were estimated using pairwise contrasts in the emmeans R package; Response, models, estimates, SE, degrees of freedom (df), z value, and p-values are reported. Significant results are in bold.

**Table S19.** Continued.

| **Within environments** | | |  | |  | |  | | |  | | |  | |
| --- | --- | --- | --- | --- | --- | --- | --- | --- | --- | --- | --- | --- | --- | --- |
| **Response** | **Environment** | **Ecotype** | | **ExT (trait) interaction, χ^2^; P-value** | | **Trend; plant size size start of season** | | **SE** | **df** | | **t. ratio** | **p-value** | | **Model** |
| seed count S1 | High environment | high | | **27.3; <.0001** | | **0.035** | | **0.006** | **470** | | **5.520** | **<.0001** | | seed count S1 ~ plant size s1*altitude+(1\|site/block) + (1\|population) |
|  |  | low | |  |  | 0.011 | | 0.028 | **470** | | 0.410 | 0.683 | |  |
|  | Low environment | high | | **146; <.0001** | | **0.110** | | **0.007** | **485** | | **16.520** | **<.0001** | | seed count S1 ~ plant size s1*altitude+(1\|site/block) |
|  |  | low | |  |  | **0.024** | | **0.005** | **485** | | **4.920** | **<.0001** | |  |
| seed count S2 | High environment | high | | **23.6; <.0001** | | **0.019** | | **0.004** | **428** | | **4.570** | **<.0001** | | seed count S2 ~ plant size s2*altitude+(1\|site/block) + (1\|population) |
|  |  | low | |  |  | -0.01 | | **0.005** | **428** | | **-1.820** | **0.07** | |  |
|  | Low environment | high | | **33.9; <.0001** | | -0.055 | | **0.020** | **385** | | **-2.7191** | **0.006** | | seed count S2 ~ plant size s2*altitude+(1\|site)+(1\|population) |
|  |  | low | |  |  | **0.014** | | **0.006** | **385** | | **2.495** | **0.0130** | |  |
| seed count S3 | High environment | high | | 2.17; 0.14 | | **0.034** | | **0.002** | **370** | | **14.360** | **<.0001** | | seed count S3 ~ plant size s3*altitude+(1\|site) +(1\|population) |
|  |  | low | |  |  | **0.042** | | **0.008** | **370** | | **5.620** | **<.0001** | |  |
|  | Low environment | high | | **53.9; <.0001** | | **0.028** | | **0.002** | **340** | | **12.930** | **<.0001** | | seed count S3 ~ plant size s3*altitude+(1\|site/block)+(1\|population/maternal family) |
|  |  | low | |  |  | **0.012** | | **0.0004** | **340** | | **28.600** | **<.0001** | |  |

| **Response** | **Environment** | **Ecotype** | **ExT (trait) interaction, χ^2^; P-value** | **Trend; plant size size start of season** | **SE** | **df** | **t. ratio** | **p-value** | **Model** |
| --- | --- | --- | --- | --- | --- | --- | --- | --- | --- |
| seed count S4 | High environment | high | **20.3; <.0001** | **0.039** | **0.001** | **314** | **34.500** | **<.0001** | seed count S4 ~ plant size s4*altitude+(1\|site/block)+(1\|population) |
|  |  | low |  | **0.018** | **0.005** | **314** | **3.800** | **0.0002** |  |
|  | Low environment | plant size s4 | NA | **0.005** | **0.001** | **185** | **4.98** | **<.0001** | seed count S4 ~ plant size s4 +(1\|site)+(1\|population) |
| seed count S5 | High environment | high | **18; <.0001** | **0.012** | **0.001** | **308** | **15.820** | **<.0001** | seed count S5 ~ plant size s5*altitude+(1\|site/block)+(1\|population) |
|  |  | low |  | **0.006** | **0.002** | **308** | **3.680** | **0.0003** |  |
|  | Low environment | high | **179; <.0001** | **0.020** | **0.001** | **181** | **15.530** | **<.0001** | seed count S5 ~ plant size s5*altitude +(1\|site/block)+(1\|population) |
|  |  | low |  | **0.004** | **0.0002** | **181** | **15.290** | **<.0001** |  |
| seed count | High environment | high | 0.15; 0.69 | **0.247** | **0.085** | **160** | **2.907** | **0.004** | mean seed ~ mean size*altitude+(1\|site/block) + (1\|population) |
|  |  | low |  | 0.139 | 0.202 | 176 | 0.687 | 0.493 |  |
|  | Low environment | high | 0.1; 0.75 | 0.145 | 0.150 | 212 | 0.998 | 0.319 | mean seed ~ mean size*altitude+(1\|site/block) + (1\|population) |
|  |  | low |  | **0.183** | **0.076** | **189** | **2.422** | **0.016** |  |

**Table S19.** Continued.

**Table S20.** Linear mixed effect models for the relationship between plant size, plant height and flowering time in the F2 populations in the low site in 2019 and high site in 2020. Significance of the trends, SE df, z values and p-values within environments for each trait. Significant results are in bold.

| **Model** | **Site** | **Trend; plant size/plant height** | **SE** | **df** | **z value** | **p-value** |
| --- | --- | --- | --- | --- | --- | --- |
| plant height 2019 ~ plant size 2019 | **Low site** | **2.496** | **0.465** | **128** | **5.362** | **<.0001** |
| flowering time 2019 ~ plant size 2019 | Low site | -0.214 | 0.111 | 126 | -1.932 | 0.056 |
| flowering time 2019 ~ plant height 2019 | **Low site** | **-0.057** | **0.011** | **128** | **-5.366** | **<.0001** |
| plant height 2020 ~ plant size 2020 | **High site** | **6.76** | **1.46** | **396** | **4.636** | **<.0001** |
| flowering time 2020 ~ plant size 2020 | **High site** | **-0.622** | **0.23** | **399** | **-2.703** | **0.007** |
| flowering time 2020 ~ plant height 2020 | **High site** | **-0.041** | **0.012** | **401** | **-3.397** | **0.001** |
